# A Single-Cell Woodchuck Liver Atlas Identifies Healthy and Disease-related Cellular Programs Conserved in Human

**DOI:** 10.1101/2025.03.03.641192

**Authors:** Zoe A. Clarke, Jawairia Atif, Xinle Wang, Lewis Y. Liu, Lawrence Wood, Damra Camat, Sai Chung, Xue-Zhong Ma, Justin Manuel, Si Lok, Timothy Lau, Cornelia Thoeni, Tomasz I. Michalak, Ian D. McGilvray, Gary D. Bader, Sonya A. MacParland

**Author notes:** equal contribution co-first authors (Zoe A. Clarke, Jawairia Atif, Xinle Wang). Equal contribution senior authors (Ian D. McGilvray, Gary D. Bader, Sonya A. MacParland). **Author Contributions:** S.A.M., G.D.B., and I.D.M. designed the study. S.L. and T.L. sequenced the genome. Z.A.C. annotated the genome. L.Y.L., D.C., S.C., X.-Z.M., L.W. and J.M. conducted liver experiments. Z.A.C. and J.A. performed the computational analysis. Z.A.C., X.W., T.I.M., and J.A. interpreted the data. C. T. performed the pathological review on the IHC. All authors contributed to the writing of the paper.

## Abstract

**Background:** Model organisms allowing for longitudinal examinations of liver disease pathogenesis are pivotal for the development of new therapeutic modalities. The eastern North American woodchuck develops chronic hepatitis and liver cancer after woodchuck hepatitis virus (WHV) infection, mirroring aspects of the natural history of the human hepatitis B virus (HBV). However, the cellular landscape of the woodchuck liver and the cell-level relevance of WHV infection to HBV infection is currently uncharacterized.

**Methods:** We employed single-cell RNA sequencing (RNA-seq) to generate an atlas of healthy woodchuck liver (63,389 cells, n=8) and peripheral blood mononuclear cells (PBMCs) (26,972 cells, n=7). Cell-specific and hepatic zonation gene signatures were validated using spatial transcriptomics (n=1). We employed our atlas to examine immune activation in stimulated precision cut-liver slices (PCLS) and disease-related pathway activation in chronic WHV infection (11,797 cells, n=3). We further employed our atlas to examine shared disease pathways between WHV infection and human HBV infection.

**Results:** Our atlas revealed woodchuck hepatic cellular diversity comparable to human and murine livers. Applying single-nucleus RNA-seq to PMA/ionomycin-stimulated precision cut liver slices revealed inflammation-associated activation signatures in T cell, myeloid and endothelial cell compartment. Finally, we describe intrahepatic T cells in chronic WHV hepatitis with both exhaustion and activation-associated signatures that resemble intrahepatic T cell genes signatures described in human chronic HBV.

**Conclusions:** We present a multi-omic atlas of healthy, diseased and *ex vivo* stimulated woodchuck liver. By identifying shared pathological processes between WHV and HBV infections, our findings reinforce the value of this preclinical model in translational research. This resource aims to advance studies on HBV pathogenesis and oncogenesis to speed the development of novel therapeutic strategies.

**Impact and Implications/Lay summary:** The liver plays important roles in metabolism, detoxification, and immune processes; liver transplantation is often the only treatment option for severe chronic liver diseases. Therefore, developing animal models that reflect human liver disease and can be studied throughout the disease course is crucial for the discovery of new treatment options. The woodchuck is an animal that develops chronic hepatitis and liver cancer after infection with woodchuck hepatitis virus, which models the human hepatitis type B virus infection (HBV) and associated hepatic carcinoma. However, our understanding of the cells that compose the woodchuck liver is limited, making it challenging to design and test cell-based therapeutics. In this study, we atlased the healthy and chronically infected woodchuck liver, found that liver cell types in woodchuck resemble those in humans, and employed the atlas to show similarities between WHV and HBV at the cell level, reinforcing the potential of WHV-infected woodchuck as a model for human HBV disease.

**Graphical abstract:** 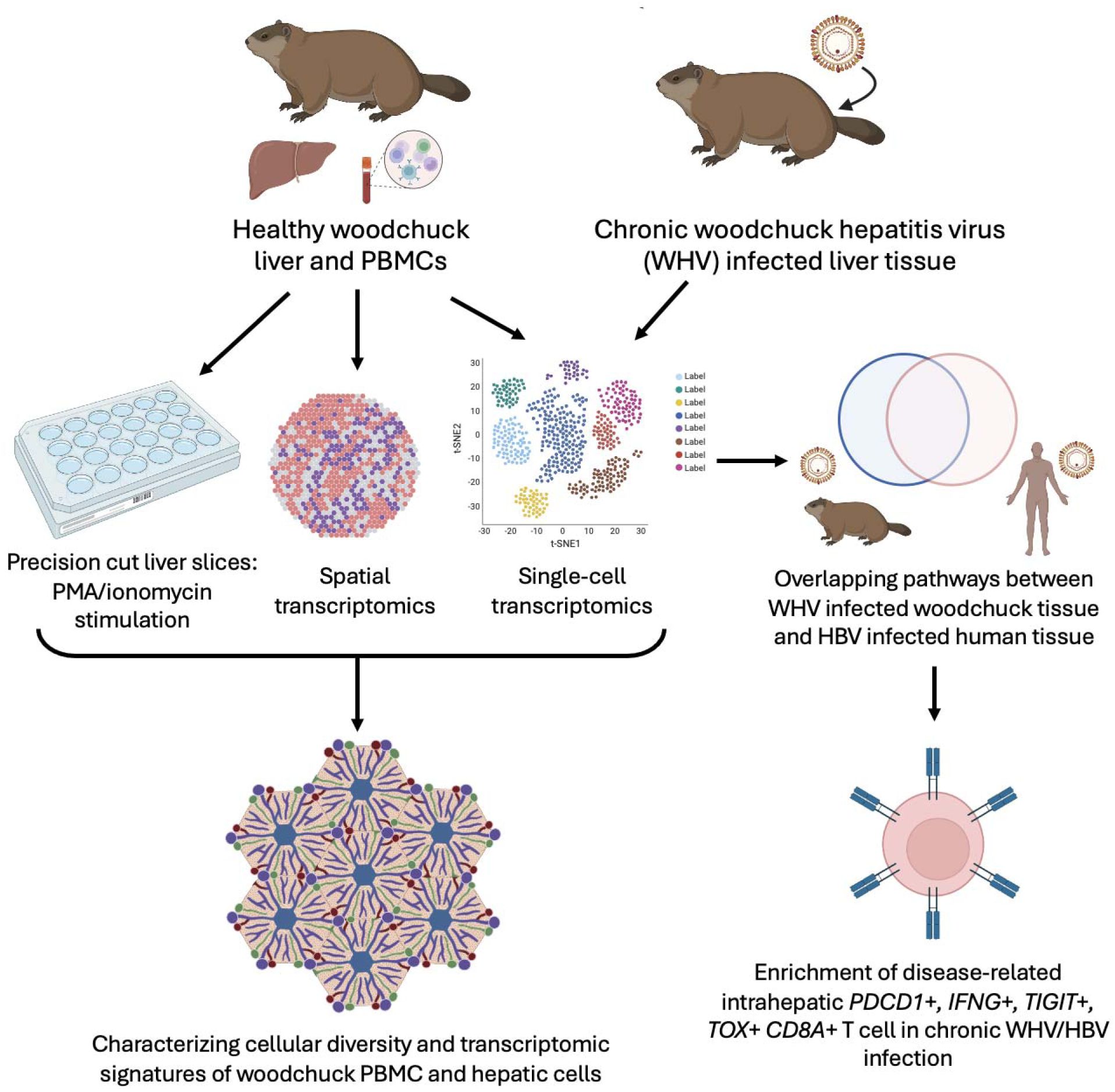

## Introduction

The mammalian liver is a vital organ with functions in metabolism, detoxification, and production of clotting factors. The liver is composed of parenchymal cell populations and non-parenchymal cell populations including tissue resident immune cells.^1^ The liver has the unique capacity to regenerate to its original size after the loss of 70% of its volume^2^. However, this regenerative capacity is abrogated in the setting of liver disease, leading to liver failure, which currently can only be treated by transplantation. Limited access to robust and physiologically relevant animal models, limited access to human liver tissue and the general fragility of certain liver cell types have made this organ difficult to study, leaving gaps in knowledge with respect to liver disease pathogenesis.^3,4^ Single cell and spatial transcriptomics has begun to narrow this gap and has broadened our understanding of the cellular complexity of the mammalian liver in health and disease, however, due to limited liver sampling as part of the standard of care in liver disease, there has been limited application of these approaches to examine the longitudinal development of disease.^5,6^

Animal models that can be serially biopsied play a critical role in longitudinal examinations of cellular ecosystems in healthy and diseased liver to enable the development of novel therapeutics. The eastern North American woodchuck, *Marmota monax*, is a preclinical animal model applied to study hepatitis B virus (HBV)-induced liver disease pathogenesis and therapeutic interventions.^7^ HBV infection was responsible for 887,000 deaths per year of which 40% were from HBV-associated liver cancer. ^8^ WHV is closely related to human HBV, with similarities in genome organization, antigenicity and immune responses induced. Furthermore, WHV infection in woodchucks closely follows the natural history of HBV and the step-by-step development of viral hepatitis finally progressing to HCC in humans^9,10^.

This model pioneered the discovery of the role of HBV in developing chronic hepatitis and primary hepatocellular carcinoma (HCC), and remains instrumental in delineating molecular immunopathogenesis of hepatitis B and evaluation of novel therapies against HBV and HCC.^11–14^ However, the full potential for this model in immunological studies remains unmet by a paucity of cell biology resources to track the cellular ecosystems in woodchuck hepatitis virus (WHV) infection, which leads to viral persistence, chronic liver inflammation, and eventual HCC presentation.

Here, we map the healthy and WHV-infected woodchuck liver at a single-cell resolution to better characterise the hepatic microenvironment in this valuable model. We annotated a new, high-quality woodchuck genome sequence to facilitate the identification of thousands of genes and transcripts by genome alignment (Fig. S1). We performed single-cell RNA sequencing (scRNA-seq) on healthy woodchuck livers and matched peripheral blood mononuclear cells (PBMCs) (Fig. 1A) and spatial transcriptomics on one healthy woodchuck liver (Fig. 1B). Finally, we expanded our analysis to assess the impact of stimulation on woodchuck intrahepatic immune responses (Fig. 1C) and applied our atlas to chronic WHV (cWHV)-infected woodchuck livers where we identified overlapping exhaustion (*TIGIT, TOX, PDCD1*) and activation (*IFNG*) pathways of T cells with T cells from liver of patients with chronic HBV infection (Fig. 1D). Our analysis of the healthy and chronically infected woodchuck livers will act as a foundation for future cell-specific studies on HBV liver pathogenesis and oncogenicity, and for pre-clinical assessments of new treatments for chronic HBV and HCC.

**Figure 1:**
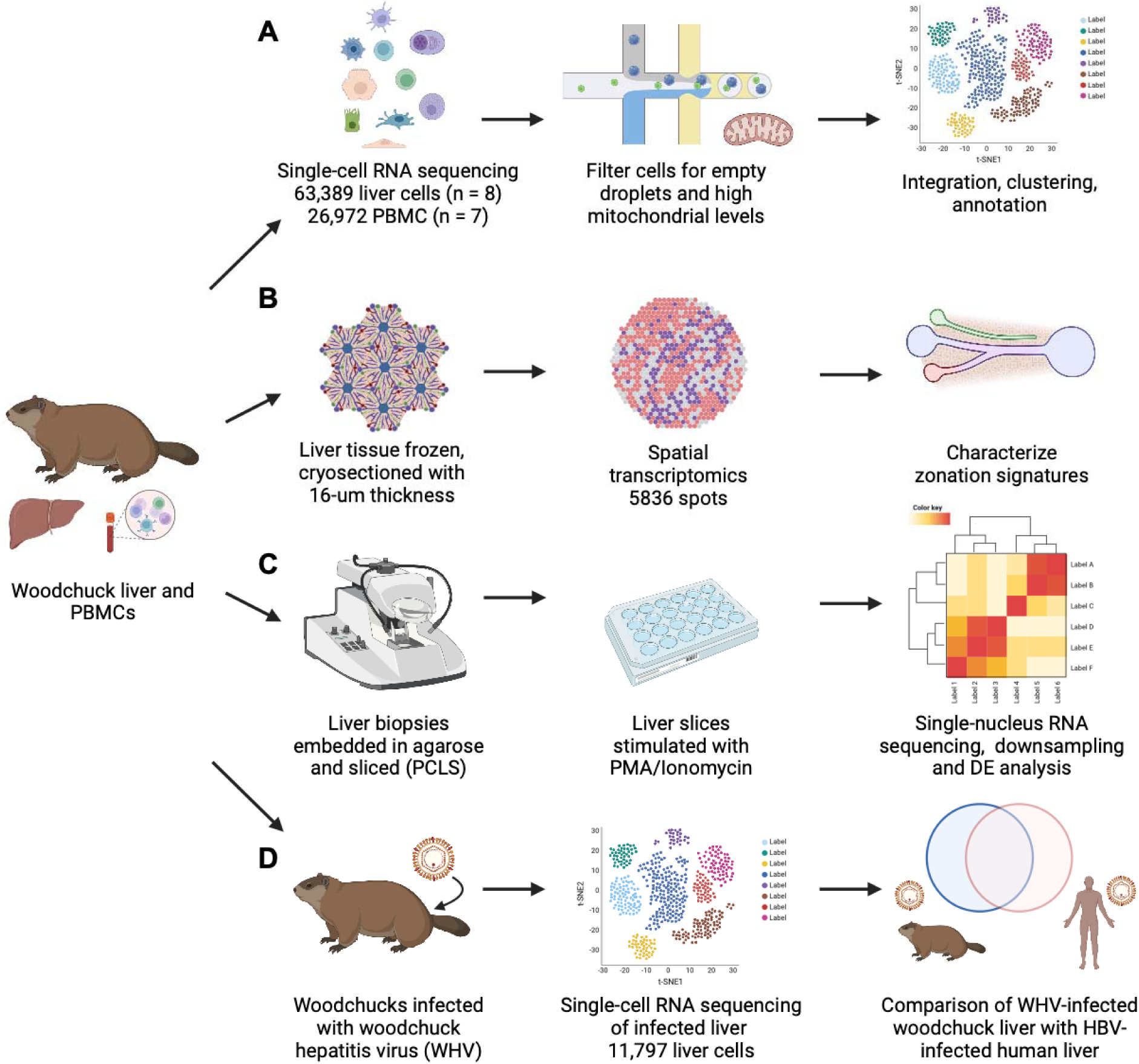
Experimental protocol for generation of the healthy woodchuck liver atlas and its application to show shared disease pathways been human HBV infection and WHV infection. The general workflow of the data processing for woodchuck liver cells, liver slices, and PBMCs. (A) Woodchuck intrahepatic cells (8 healthy livers) and PBMCs (7 matched blood samples) were isolated and underwent single-cell RNA sequencing with 10X Genomics. Cells with mitochondrial content levels past a threshold were then filtered out, and cells had to be classified as cells rather than empty droplets using DropletQC. The cells were then clustered, integrated, and annotated using the Seurat pipeline and manual annotation. (B) For the spatial transcriptomics analysis, serial liver slices from 1 healthy woodchuck were processed following the protocol outlined in Andrews *et al.* (2022).^26^ The data were then analyzed with 10X Genomics’ spatial transcriptomics pipeline, and zonation signatures were characterized by identifying differentially expressed genes with Seurat. (C) For the stimulation experiments, precision cut liver slices (PCLS) were stimulated with PMA/Ionomycin, and the stimulated data were downsampled to create a comparable sample size as the unstimulated data. (D) Livers were harvested from woodchucks chronically infected with WHV and scRNA-seq was performed with cells filtered as in (A). Cells from diseased livers (n=3) were merged with healthy woodchuck liver cells (n=2) and compared to human cells obtained from HBV- infected livers (n=5) and healthy human liver controls (n=6).

## Methods

### WHV infection, liver tissue processing and PBMC isolation

Samples were collected from eight healthy female outbred woodchucks and three WHV-infected outbred woodchucks (woodchuck characteristics described in Table S1). Prior to tissue dissociation, a small section of each sample was formalin-fixed, paraffin embedded and sectioned for haematoxylin and eosin staining. For enzymatic perfusion, woodchuck livers were procured from Memorial University of Newfoundland, St. John’s, NL, Canada, under the approval of the Institutional Committee on Animal Bioethics and Care (protocol 15-155-M). Three woodchucks were infected with a single dose of strain WHV7 inoculum with 6.5 x 10^10^ DNase digestion-protected virus copies, equivalents of the virion genome. Tumor-distal WHV infected samples were collected at the time of progression to HCC (as detected by GGT elevation>50 U/L).Two healthy and the three WHV-infected woodchuck livers were dissociated according to a previously described 2-step collagenase perfusion protocol (supplementary methods)^5^. Healthy woodchuck liver biopsies (n=6) were conducted under approval of the Animal Resources Centre at the University Health Network, Toronto, ON, Canada (AUP 6199.6). Woodchucks were sedated as described in the supplementary methods, and liver biopsies were obtained with 8-mm biopsy forceps through incision sites of approximately 1.5 cm along the midline 1.5-2.0 cm below the diaphragm. The tissue was dissociated for 30 minutes at 37 in 0.2mL of pre-warmed mixture of 0.275 unit of neutral protease (VitaCyte, cat. 003-1000) and 2.5×10^4^ units of a collagenase mixture of 60% class I and 40% class II collagenase (VitaCyte, cat. 001-2030) in 1x of Hank’s balanced salt solutions, as described previously.^15^

Whole blood was collected in EDTA tubes. Plasma was collected via 1000 x g spin for 10 minutes. PBMCs were isolated via a Ficoll density gradient. The resulting PBMCs and tissue homogenates were strained through a 70-µm cell strainer on ice and sent for scRNA-seq processing at the Genomics Core of the Princess Margaret Cancer Research Tower in Toronto, ON, Canada.

### Woodchuck precision cut liver slice culture

Woodchuck precision cut liver slices (PCLS) were generated and cultured as previously described.^16^ Briefly, 3-mm punch biopsies (Integra Miltex) were embedded in agarose (see supplementary methods). Slices of 250-μm thickness were obtained using a Leica VT1200S Vibratome machine. One slice per well was plated in 24-well plates containing 500 μL of culture medium and organotypic cell inserts (Sigma Aldrich PICM01250). The plates were incubated overnight on an orbital shaker at 18 RPM before being either stimulated with phorbol myristate acetate (PMA) (100 ng/mL) and ionomycin (1 μg/mL) or kept unstimulated as a control. Slices were collected after 6 hours of stimulation and snRNA-seq was performed on four slices per condition, as described below.

### Single-cell and single-nucleus RNA-sequencing and data processing

For both liver tissue and PBMC datasets, cells remained unsorted due to both the fragility of hepatocytes and cholangiocytes.^5^ Single-cell samples were sequenced with 10X Genomics SingleCell 3’ v2 chemistry^17^ and were sequenced on a HiSeq 2500 or NovaSeq 6000.

For woodchuck PCLS experiments that were examined by single nucleus RNA sequencing, nuclei were extracted from 4 slices per condition using a CHAPS with salts and Tris (CST) protocol, as previously described (see supplementary methods). ^18^ Nuclei were resuspended, stained with a 1X salts and tris solution containing DAPI, and sorted using fluorescence-activated cell sorting with an ARIA II FACS sorter.

To support the annotations of our single cell data, we generated and annotated a high-quality woodchuck genome with 64.6-94.8% of reads confidently mapped to the genome and 41.3-77.6% confidently aligned to the transcriptome (Table S1, Supplementary Methods: Mapping and annotating a reference woodchuck whole genome). The closest published assembly to this genome is GCA_021218885.2 (link to genome: https://doi.org/10.5281/zenodo.10855128).

Sequenced single-cell and single-nucleus reads were aligned to the woodchuck genome with 10X Genomics Cell Ranger v5.0.0, and filtered reads were processed with the Seurat pipeline (v5.0.0)^19^. The percentage of mitochondrial genes per cell were calculated for all samples. For single-cell data, cells with greater than 10% of mitochondrial content were removed from downstream analysis of PBMCs, and cells with greater than 50% mitochondrial content were removed from liver tissue as previously described.^5^ Mitochondrial genes were removed from the gene-by-cell matrix before clustering. Scaling and normalisation were done with Seurat’s SCTransform pipeline. Empty droplets were predicted and removed with DropletQC under default settings for single-cell liver tissue (quality control steps summarized in Fig. S2A-G). Hepatocytes were removed from liver biopsy samples, because the gene expression profiles for these cells appeared corrupted and inconsistent, likely attributed to being negatively impacted by the protocol. The PCLS-stimulated dataset was randomly downsampled to create even populations of stimulated and unstimulated data.

Marker genes for cell clusters were interpreted at different clustering resolutions with scClustViz, a single-cell transcriptomic map visualisation tool.^20^ Seurat’s Canonical Correlation Analysis (CCA) function, a tool that merges single-cell datasets and applies batch correction across samples, was applied separately to healthy PBMCs, healthy liver tissue homogenates, and PCLS-stimulated single-nucleus liver tissue with default parameters. In addition, diseased single-cell liver tissue was merged with the healthy liver tissue homogenate single-cell data with no integration to avoid over-integration. This combined all cells across each respective data type, creating two healthy single-cell maps of integrated samples (i.e. maps of both woodchuck PBMCs and tissue liver homogenate), one map of PCLS-stimulated single-nucleus liver tissue, and one healthy and diseased single-cell map of liver tissue homogenate single-cell data.

For the healthy and PCLS-stimulated datasets, the batch-corrected data from the integrated maps were used to cluster the cells, but the gene expression data normalised with SCTransform were used for differential expression testing, as recommended by Seurat documentation.^21^ Principal component analysis (PCA) was performed using Seurat’s default parameter of 50 principal components (PCs), then nearest neighbours were calculated and UMAP clustering performed using the first 30 PCs. Graph-based clustering was performed at different resolutions that best represented cell heterogeneity, the resolutions were as follows: 0.6 for PBMCs, 1.4 for healthy liver tissue, 0.4 for the PCLS-stimulated single-nucleus liver tissue, and 1.6 for the merged healthy and cWHV-infected liver tissue. Clusters were manually assigned the same label to create cell populations composed of multiple clusters. For the merged healthy and cWHV-infected liver tissue maps, the PrepSCTFindMarkers function was run to prepare the map for differential gene expression (DGE) analysis since multiple SCT models were being combined. Presto (v1.0.0) was used to calculate maker genes, retaining only positively altered markers that meet two criteria: expression of at least 25% of the cells in a cluster (min.pct=0.25), and a minimum log2 fold change in expression of 0.25 (logfc.threshold=0.25).^22^ Clusters were annotated based on top differentially expressed genes and lineage-associated marker expression. For pairwise healthy vs cWHV-infected DGE analysis per cell type, Seurat’s^21^ FindAllMarkers function was used with a min.pct=0.15 and a logfc.threshold of 0.1. Most plots were generated with Seurat (v5.0.0), ggplot2 (v3.4.4)^23^, dplyr (v1.1.4), and rcartocolor(https://github.com/Nowosad/rcartocolor), and scCustomize (v1.1.3)^24^. EnhancedVolcano (v1.22.0)^25^ was used to generate Figure 5F from these results.

### Spatial transcriptomics

Spatial transcriptomics was performed on frozen OCT embedded healthy (sample L212) woodchuck liver tissue sectioned to a thickness of 16-μm at −10 following the protocol described in *Andrews et al. (2022)*^26^. The data was analysed with 10X Genomics Space Ranger v1.2.2 and processed with Seurat’s v4 spatial transcriptomics analysis pipeline.^19^ The SCTransform normalisation method was performed separately on two liver slices, which then had their variable features combined for downstream analysis, as recommended by Seurat (Seurat: vignettes/spatial_vignette.Rmd). From these combined features, dimensionality reduction with 50 PCs and nearest neighbour and UMAP reduction with 30 PCs was performed. Clustering was done at 0.2 resolution and DGE analysis was carried out for marker gene and pathway analysis using Seurat’s FindAllMarkers function (min.pct=0.25 and logfc.threshold=0.25, only.pos=TRUE). After assigning periportal and pericentral hepatocyte identities to two clusters, an additional comparison of periportal and periportal clusters was performed using Seurat’s FindMarkers function with the same parameters to find genes that best distinguished the two cell populations.

### Pathway analysis

Pathway analysis was performed by running Gene Set Enrichment Analysis (GSEA)^27^ on ranked lists of differentially expressed genes from cluster-cluster comparisons. Results from GSEA were visualised with the EnrichmentMap v3.3.1 and AutoAnnotate v1.3.3 tools from Cytoscape v3.8.2^28^. The gene sets used as input for GSEA were from a custom file containing woodchuck gene sets that were converted from a human gene set file (Human_GO_AllPathways_no_GO_iea_November_01_2021_symbol.gmt) using human-woodchuck orthology that we defined in our genome annotation pipeline (Fig. S1).

### Woodchuck-human comparison for healthy liver

Well-established marker genes used to identify cell types in mice and humans from resources such as the Human Protein Atlas^29^ and GeneCards^30^, were used to identify general woodchuck cell identities (e.g., *CD3E* for T-cells and *MARCO* for liver tissue resident macrophages). After general cell identities were assigned, top marker genes were examined in the literature to interpret and identify cell subtypes, and human or mouse cell subtype markers were also searched for in the woodchuck data. The gene expression profiles of the liver clusters were compared to existing liver gene expression data^5,26^ using one-to-one orthology mapping and Spearman correlations (Fig. S3A-D).

### cWHV-infected woodchuck comparison to HBV-infected human tissue

To examine if our woodchuck atlas could be applied to as the question of whether cWHV and HBV infections result in shared disease-related immune dysfunction profile, we sub-clustered T cells from a publicly available scRNAseq dataset generated from biopsies taken from chronically HBV infected individuals in the immune-active phase of infection (n=5) with 6 healthy liver controls from the same dataset. Thus, count matrices from published HBV infected human liver data (GSE182159)^31^, were merged and then normalised and scaled using SCTransform in Seurat. Data did not require batch correction as samples merged well without further transformation and PCA was performed to calculate 30 components, all of which were used to construct a shared nearest neighbour graph and generate UMAP embeddings. A resolution of 1.2 was used for clustering with the Louvain algorithm. Clusters were annotated based on lineage-specific markers, as described in the supplemental materials and the original publication. The R package fgsea (v1.30.0)^32^ was used to identify significantly enriched pathways in clusters of both cWHV-infected woodchuck (determined using the human-woodchuck orthology based gene set generated above, *p*-value <0.1) and HBV-infected human liver samples (determined using the human gene set file listed above, *p*-value <0.1) relative to their respective healthy controls and pathways significant in both cases were plotted using ggplot2 (v.3.5.1) and ComplexHeatmap (v2.20.0).^33^

## Results

### An atlas of healthy woodchuck liver tissue

Combining the scRNA-seq liver tissue data from six core biopsies and two perfused caudate liver lobes (Fig. 2B, Fig. S2H), 213,090 liver cells were detected by Cell Ranger before additional filtering parameters were applied (Table S1), which was reduced to 100,595 cells after mitochondrial content cutoff (50%)^5^ and a minimum of 200 features per cell criteria were applied. Most cells removed by the mitochondrial cutoff were pericentral hepatocytes, and further lowering of the cutoff threshold did not significantly impact the results (Fig. S2G). Subsequent filtering of empty droplets with DropletQC^34^ and removal of biopsy hepatocytes resulted in 63,389 cells (liver map Fig 2A,E, DropletQC filtering Fig. S2A-F). Cells removed by DropletQC tended to demonstrate strong batch effects and expressed high levels of hepatocyte background genes (e.g. *APOC1, APOA1*), suggesting that these cells were either empty droplets or highly contaminated containing fragments from multiple cell types (Fig. S2E,F). After filtering with DropletQC, cell types consistent with those found in the human liver were described (marker gene and cell lineage summaries Fig. 2G,I and Table S2, S3).

**Figure 2:**
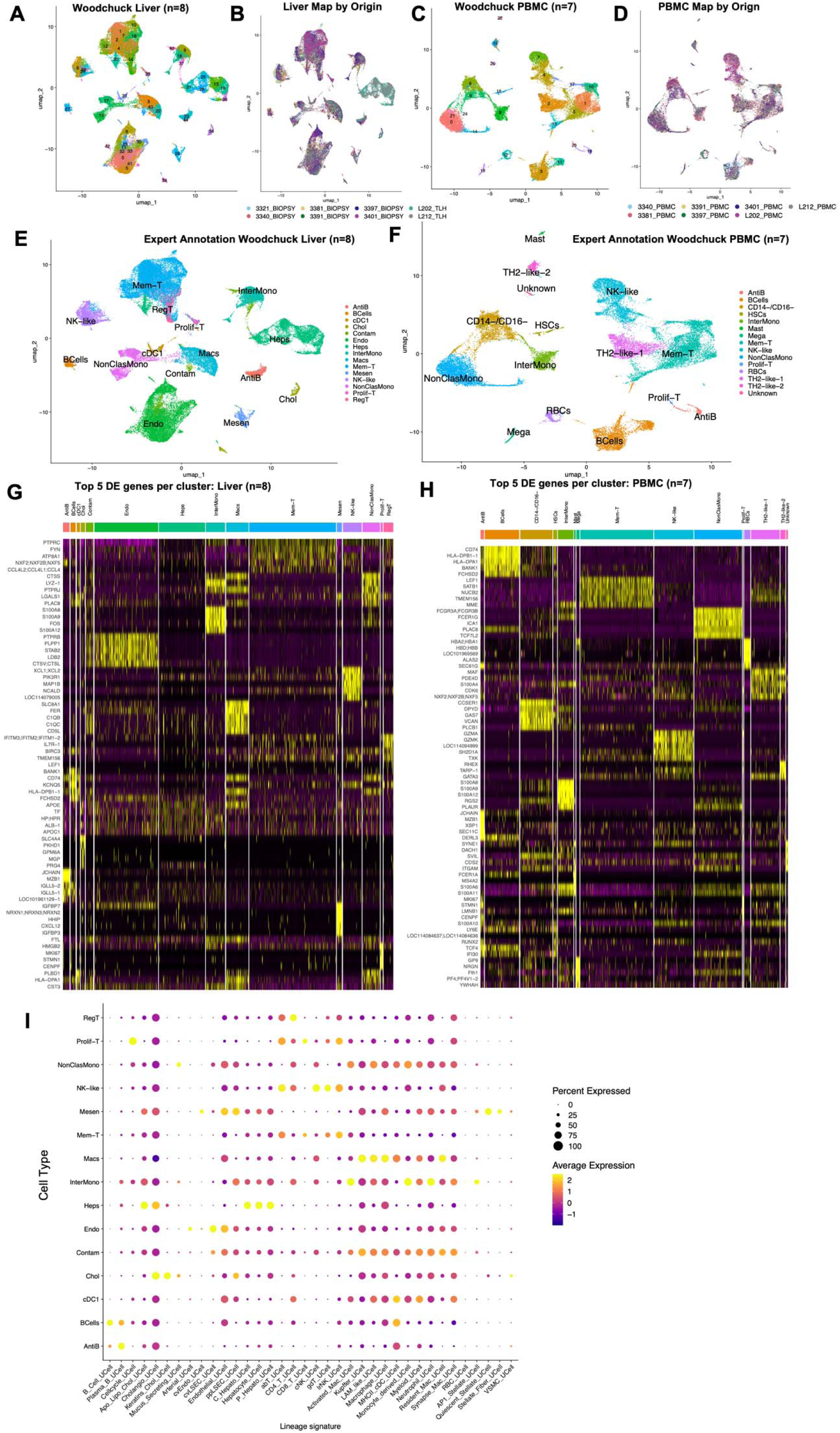
Woodchuck liver and PBMCs show distinct populations of circulating and tissue-resident immune cells. (A) A UMAP of 63,389 cells from woodchuck liver biopsies (n=6) or liver homogenate from perfused caudate lobes (n=2) labelled by cluster number.(B) Woodchuck liver cells grouped by sample identity on a UMAP. (C) A UMAP of 26,972 woodchuck PBMC from n=7 woodchucks, also labelled by cluster number. (D) Woodchuck PBMCs grouped by sample identity on a UMAP. (E) A UMAP of woodchuck liver cells and (F) PBMCs with their assigned cell identity in short notation.(G) A heatmap of top 5 differentially expressed genes for each cell identity in the woodchuck liver. (H) A heatmap of top 5 differentially expressed genes for each cell identity in woodchuck PBMCs. (I) Previously identified marker genes for different liver cell populations^39^ were used to score cell lineage signatures with UCell (v2.4.0)^86^ to the different woodchuck liver cell types presented as a dot plot. Lighter colours indicate higher expression values, and larger dots indicate that a larger percentage of cells in the group express that signature.

PBMC scRNA-seq data were collected from seven woodchucks (Fig. 2D, Fig. S2I). Samples were processed with default filtering settings by Cell Ranger and filtered with standard 10% mitochondrial content cutoff, which left 26,972 total PBMC cells (Fig. 2C,F). Subtypes consistent with known PBMC heterogeneity were detected (marker gene summaries Fig 2H and Table S4, S5).

Unbiased spatial transcriptomics (Visium) was applied to ascribe a geographical pattern of the different cell types in the woodchuck liver. This was able to provide spatial context to periportal and pericentral hepatocytes (Fig 3A-E). However, we were unable to discern the spatial patterns of other cell types due to the dominant gene signatures provided by the hepatocytes. Taken together, this atlas provides a platform for examining the cell biology of the woodchuck liver and annotating the intrahepatic cell populations (described below).

**Figure 3:**
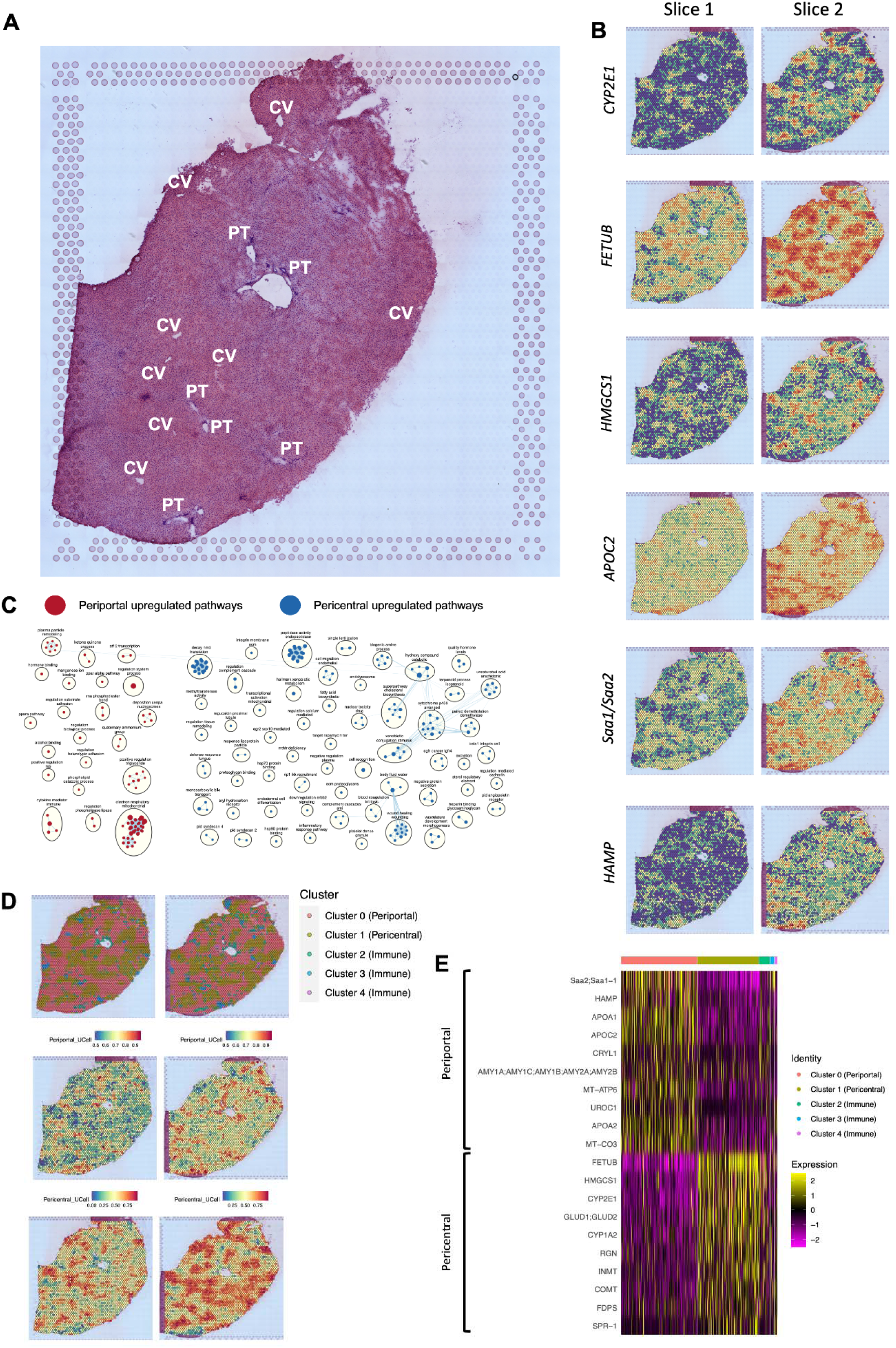
Spatial transcriptomics data describes zonation patterns in the woodchuck liver. Spatial transcriptomics was applied to 2 slices of healthy woodchuck liver. (A) H&E stained woodchuck liver annotated for zonation by a pathologist. Zones were labeled as central venous/pericentral (CV) or periportal (PP) depending on their pathology. (B) Top pericentral *(CYP2E1, FETUB, HMGCS1)* and periportal *(APOC2, Saa1;Saa2, HAMP)* differentially expressed genes identified with their expression projected onto two liver slices (Slice 1 and Slice 2) as a heatmap. Blue colours indicate low expression, and red colours indicate high expression. (C) Pathway enrichment analysis was performed on periportal and pericentral DE genes. Pathways were identified with GSEA and visualized with Cytoscape. Each dot (node) represents a pathway, with the size of the dot indicating the number of genes in the pathway. Related pathways are connected by light blue lines, grouped together in black circles, and labelled. (D) Spatial transcriptomics clusters projected onto the liver slice (top), and periportal (middle) and pericentral (bottom) gene signatures applied to spatial transcriptomics liver data with UCell^86^. Signatures were created using the top 10 pericentral and periportal genes as determined by clustering and differential gene expression of the spatial transcriptomics data (Table S7), and signature scores are projected as a heatmap onto the slices. (E) A heatmap of the top 10 periportal and pericentral genes grouped by clustering of the spatial transcriptomics data. Periportal and pericentral hepatocytes are represented by clusters 0 and 1 respectively. Clusters 3 and 4 did not represent any liver zonation.

### Hepatocytes

Hepatocytes have distinct gene expression profiles based on their proximity to the portal triad and central vein (zones labeled on a liver slice by a pathologist Fig 3A).^35^ Periportal hepatocytes are more oxygenated than pericentral hepatocytes, allowing them to possess energy expenditure behaviours like cholesterol biosynthesis and oxidative metabolism.^36^ Conversely, pericentral hepatocytes specialise in lower energy behaviours, such as glycolysis and xenobiotic metabolism.^36^

Spatial transcriptomics confirmed two distinct pericentral and periportal hepatocyte phenotypes in our woodchuck samples (clusters 1 and 0 respectively, Fig 3B-E, Tables S6, S7, and S8). The pericentral hepatocytes identified in our samples shared well-characterised pericentral hepatocyte markers with the human liver such as *CYP2E1*, *FETUB*, and *HMGCS1* (Fig. 3B), as well as other top identified markers including *GLUD1/GLUD2, CYP1A2, RGN, INMT, COMT, FDPS,* and *IDI1* (Table S6, S7). Interestingly, *HMGCS1* in our sample was expressed in the pericentral regions despite it being previously identified as a periportal marker in some human studies.^5,26^ From our pathway analyses, we found that enriched pathways in the pericentral zones include those of xenobiotic metabolism, blood coagulation, bile transport, and demethylase activity. Electron respiratory chain pathways were also decreased, which is consistent with the human pericentral phenotype (Fig. 3C).^35^

Periportal hepatocytes in the woodchuck shared both human periportal markers *HAMP, APOC2,* and mouse orthologs *Saa1/Saa2* (Fig 3B), in addition to upregulated markers *APOC3, TTR, CRYL1, APOA1, UROC1, APOA2* and an ortholog to the *AMY1/AMY2* genes (Table S6, S7). However, in the scRNA-seq data, these cells were mainly identified by the lack of pericentral markers, while periportal markers were a source of ambient RNA contamination (Table S3). An exception to this was *PCK1*, which was clearly expressed in a small proportion of periportal hepatocytes, suggesting that these cells may be damaged or otherwise influenced by the experimental protocol, thus making them difficult to identify (Fig. S4). Pathway enrichment analysis of periportal zones from the spatial transcriptomics data revealed triglyceride regulation, alcohol binding, hormone binding, and electron respiratory chain pathway activity (Fig. 3C).

A third subset of hepatocytes was also identified from the scRNA-seq data, which did not clearly represent a periportal or pericentral phenotype. Some of the top differentially expressed genes of this population include *C8G, LOC114082496* (cytochrome P450 2C19-like)*, LOC106144883* (translation initiation factor IF-2-like)*, GADD45G,* and *SLC22A11* (cluster 36, Fig 2A, Table S3). These cells are clearly distinguished by their high nuclear fraction values compared to all other hepatocytes, showing that they have the highest proportion of unspliced transcripts (mean of 50.4% unspliced, compared to total hepatocyte average of 21.5%) across all single-cell clusters. Top pathways associated with this cluster when compared to other hepatocytes are related to synapses, G protein-coupled receptors, and cytoskeleton regulation; negatively regulated pathways include coagulation, the complement cascade, and translation. However, it is unclear whether this is a biological signal, or a technical artefact related to the scRNA-seq tissue dissociation protocol, given that these patterns were not present in the spatial transcriptomics data.

Overall, our examination of hepatocytes in woodchucks show clear relationships with human hepatocytes, especially when comparing top zonation markers from spatial transcriptomics. Although the relatively lower quality of hepatocytes in single-cell data presents a challenge, the shared cross-species patterns found in our analysis reinforces the woodchuck as an appropriate model for the human liver.

### Endothelial cells

Endothelial cells constitute a large proportion of hepatic cells and are involved in the maintenance of liver homeostasis, regulating quiescence of immune players such as Hepatic stellate cells (HSCs) and Kupffer cells^37,38^. Previously, we identified three populations of endothelial cells in the human liver associated with location: periportal liver sinusoidal endothelial cells (LSECs) (Zone1), central venous LSECs (Zone 2 or 3), and a small population of non-LSEC endothelial cells.^39^ These populations were distinguished by an expression of *CALCRL* and *RAMP2*. In the woodchuck liver map, we identified a cell population (n = 12,741) with the conserved expression of these two shared endothelial markers, along with additional LSEC markers *CD32B (FCGR2B), LYVE-1, and STAB2*^5^ (endothelial cell cluster in Fig. 2E, markers visualized in Fig S5).

This endothelial-like cell population also showed a strong expression of *CLEC4G* which is characteristic of LSECs (Fig S5).^40^ We speculate that different types of endothelial cells exist within this large LSEC cluster, possibly including a mixture of hepatic artery endothelial cells based on *CD34, MECOM, VWF,* and *BMX* expression and lower *CLEC4G* expression^40–44^; and central venous endothelial cells based on *RSPO3, WNT2* and *ACKR1* expression^42,45^ (Fig. S5, Table S2). These two populations are on opposite sides of the LSEC cluster, and their respective marker genes decrease in expression towards the centre of the cluster, which likely consists largely of LSECs lining the sinusoid connecting the central vein and hepatic artery (Fig. S5). The expression gradients and lack of distinct clustering suggests that LSEC subpopulations are not distinct, but rather exist on a phenotypic gradient.

Low expression of these genes observed in the spatial transcriptomics data prevented confirmation of the zonation of these markers. However, presence of these zonated populations may be confirmed in future spatial experiments with higher cellular resolution.

### Mesenchymal cells

Mesenchymal cells in the liver encompass cell populations of central venous and periportal fibroblasts, vascular smooth muscle cells, and HSCs, which are the major collagen-producing cells.^46,47^ We identified Cluster 26 (n = 1,069) in the woodchuck liver map to be mesenchymal-like (Fig. 2A,E) with its expression of many genes differentially expressed in human HSCs, such as *COL1A2, COL3A1, IGFBP7, IGFBP3, DCN*, *CALCRL, COL1A1, SPARC, RBP1,* and *HHIP* (marker genes found in Fig. S6, Table S2).^5,26^ Interestingly, mesenchymal cells in the woodchuck appear to have the most conserved human differential gene expression compared to all other cell types.

Mesenchymal cell populations between humans and woodchucks have strong Spearman correlation coefficients when comparing gene expression data (Fig. S3A,B,D). Furthermore, many top mesenchymal cell marker genes are conserved in the human liver.^5^ Quiescent mesenchymal cell markers, *LRAT* and *PDE3B,* and activated mesenchymal cell markers, *ACTA2, SPARC,* and *COL1A1*, however, largely overlap, suggesting that it is difficult to resolve distinct mesenchymal cell subtypes in this dataset.^26^ Even though subtypes are indistinguishable in this dataset, we can clearly see that mesenchymal cells and many of their distinguishing marker genes are conserved between woodchuck and human.

### Cholangiocytes

Cholangiocytes are liver cells located in the bile duct and responsible for the creation of bile. Each bile duct is enclosed by 4-12 cholangiocytes depending on the size of the bile duct. Common markers that characterise cholangiocytes include *KRT19, EPCAM, SRY,* and *SOX9.*^5,26,40^ Full transcriptomic assessment of human cholangiocytes are scarce due to the difficulty in isolating these cells. We previously identified the transcriptome of cholangiocytes in five human livers with our dissociation protocol.^5^ Using the same dissociation protocol in the woodchuck livers, we uncovered two populations of cells resembling cholangiocytes (n = 810, clusters 34 and 38, Fig. 2A,E), which express genes differentially expressed in human cholangiocytes, including *ANXA2*, *CST3*, *BIRC3*, *TESC, and KRT19* (markers found in Fig. S7, Table S2).^5,26,40^ Cluster 34 also expressed *SOX9* and *EPCAM,* while cluster 38 did not. Cluster 38 expressed mesenchymal and endothelial cell markers suggesting that this may be a cluster of doublets, while cluster 34 seemingly represents a true cholangiocyte population. Low expression of cholangiocyte markers in the spatial transcriptomics data prevented an estimation of zonation from being applied to the cells. However, the conservation of cholangiocyte markers and high correlation coefficients between human and woodchuck cell types (Fig. S3A,B,D) supports a consistent cholangiocyte phenotype across both species.

### T cells

CD3D and CD3E expression, as well as the expression of GIMAP genes, including *GIMAP1, GIMAP4, GIMAP5, GIMAP6,* and *GIMAP7*, largely associated with T cell function, were seen in 23,387 cells of the woodchuck liver map, clustering to four distinct populations.^29,48^ (Fig. 2A,E). We identified populations of regulatory T cells (cluster 20, n = 1866), NK/T cells (clusters 6, 24, 29, n = 3804), tissue-resident memory T cells (clusters 1, 2, 4, 7, 10, 12, 16, n = 17,305), and proliferating T cells (cluster 35, n = 412) based on known human T cell markers (Fig 2A,E, marker genes found in Fig. S8, and Table S2).

The population resembling regulatory T cells (cluster 20) was distinguished by regulatory T cell markers such as *CD4, CD28, FOXP1, CTLA4,* with *CD38* (a marker of recent T cell activation)^49^ and *CD44* (activation marker)^50^ NK-like clusters expressed *XCL1/XCL2, IL2RA (CD25), CD38,* and *NKG7* (Fig. 2A,E, Fig. S8, Table S2).^51^ Meanwhile, for clusters 1, 2, 4, 7, 10, 12, and 16, *CD8A, CCL5, CD69, CD2,* and *GZMK,* but not *CD25* or *CD38* expression in the tissue-resident memory clusters (Fig. 2A,E, Fig. S8, Table S2) suggest that these are likely not newly activated cells and may in fact be a resting tissue-resident memory T cell population.^51,52^ Finally, cluster 35 expressed general T-cell markers in addition to *BRCA2, TOP2A,* and *STMN1,* which are proliferating markers, suggesting these are cycling T cells (Fig. 2A,E, Fig. S8, Table S2).

In the PBMC dataset, we observed five distinct T and NK-like populations (Fig. 2C,F) including NK-like cells (cluster 4 and 7, n = 3,580), memory CD4+ T cells (clusters 1, 5, 10, 13, 17, n = 6,628), proliferating T cells (cluster 12, n = 77), and two populations of TH2-like T cells (population 1, cluster 2, n = 2,606; population 2, cluster 15, n = 452; Fig. 2C,F, marker genes found in Table S4, S5). The population of NK-like cells expressed *XCL1;XCL2* and *NKG7* (Fig. S9, Table S4), while the population of memory CD4+ T cells expressed markers *LEF1, IL7R, CD4, TCF7, CCR7*, *CD28, PDCD4, CD38, and ZEB1* which suggests a mature, resting population (Fig. S9, Table S4).^51,53,54^ Proliferating T cells expressed markers *BRCA2* and *STMN1,* while the expression of markers *GATA3, MAF, CCR4,* and *S100A4* support the TH2-like nature of the remaining two cell populations^55–57^, with one of the populations (cluster 2) expressing more *CXCR3* and *IL2RA (CD25)* and the other (cluster 15) expressing more *CD160* (Fig. S9, Table S4).

### B cells

Two clusters in our liver map were identified as B cell populations based on their marker expression (Fig. 2A,E). Clusters 27 (n = 1063) and 23 (n = 1,291) share markers associated with the B cell signature, including *CD19, MS4A1 (CD20), CD79A, CD74* and *IGLL5*^58^, while cluster 27 additionally expressed *JCHAIN* and *MZB1*, characteristic of plasma cell differentiation, suggesting this may be a population of antibody-secreting B cells (Table S2).^59,60^ Two populations of B cells were likewise identified in the PBMC dataset based on similar gene signatures: cluster 3 and 11 (n = 3,132) and cluster 18 (n = 306), resembling B cell and plasma B cells, respectively (Fig. C,F, Table S4).

### Myeloid cells

In the human liver, we have previously identified multiple myeloid populations that include a spectrum of recently recruited monocyte-like cells to tissue resident macrophages ^5^ In the woodchuck liver, all myeloid cells expressed DE genes similar to those shown in human liver monocyte/macrophage populations, including *TYROBP, CD74* and *CTSS*^5,26^. We found four myeloid clusters, with three of these populations suggesting a more monocyte-like phenotype, and one with a more tissue resident phenotype. The more monocytic-like populations resembled CD14+/CD16+ intermediate monocytes (n = 3,760), CD14-/CD16+ non-classical monocytes (n = 3,500), and cDC1 conventional DC type 1 cells (n = 633), whereas the tolerogenic population were labelled as macrophages (n = 4,468) (Fig. 2A,E).

In addition to *TYROBP, CD74* and *CTSS,* all myeloid populations expressed distinct marker genes which guided their annotation. For instance, clusters 9 and 18 (Fig 2A, n = 3,760) expressed strong signals of CD14, CD16*/FCGR3A, S100A8, ITGAM/CD11b, SIRPA, NOTCH2, IL17RA* and *LYZ* (Table S3), suggesting resemblance to CD14^+^/CD16^+^ monocytes.^61^ The second myeloid population (clusters 15 and 17; Fig 2A, n = 3,500) shared the same markers, but with stronger CD74 and CD16, and weaker CD14 expression, analogous to CD14^-^/CD16^+^ non-classical monocytes (Table S3).^61^ The third population (cluster 31, Fig 2A, n = 633) expressed markers associated with conventional DC type 1 (cDC1) cells including *BTLA, CADM1, ID2, LY6E, FLT3* and low levels of *SIRPA* (Table S3).^62^ The final population (clusters 3, 22 and 43, Fig 2A, n = 4,468) is distinguished by many human Kupffer cell markers, including *MARCO, HMOX1, CD5L, VSIG4*, *VCAM1* and *CD68* (Table S3).^5^ Genes encoding the complement proteins were also among the top differentially expressed genes.

In examining circulating woodchuck PBMCs, we found similar populations of CD14+/CD16+ intermediate monocytes (n =1,430 cells), CD14-/CD16+ non-classical monocytes (n = 4,326 cells), and a new population of CD14^lo^/CD16-monocytes (n = 2,941 cells), resembling classical monocytes which represent a large portion of circulating monocytes^63^. This PBMC dataset also contained a population resembling basophils or mast cells (n = 125 cells) with DE genes *FCER1A* and *MS4A2* (Fig. 2F, Table S4). These genes are involved in the allergic response and are expressed on the surface of mast cells and basophils.^29,64,65^ This population also expressed *Prss34*, an ortholog to a mouse gene that encodes mast cell protease found in both cell types (Table S4).^66^

### Examination of woodchuck intrahepatic cell function using precision-cut liver slices

To support our cell cluster annotations and determine woodchuck cell-specific responses to inflammatory stimuli, woodchuck precision-cut liver slice (PCLS) samples were stimulated *in vitro* with PMA and ionomycin before characterization using snRNA-seq (Fig 1C, Fig. 4A). This approach leads to receptor ligation-independent PMA induced NF-kB pathway activation and ionomycin-induced increase in intracellular calcium in tissue resident cells.^67–69^ Cell types were identified based on their gene expression profiles (Fig. 4B, Table S9), and PMA-stimulated cell populations were directly compared to their unstimulated counterparts on a per-cluster basis (Fig. 4C-E, Table S10). PMA-stimulated myeloid populations were characterised by a higher expression of *NFKB1, TNF, IFNG, IL2RA, IL7, CCL4,* and *CCL3* (Fig. 4F,G), which are markers associated with immune activation. Pathway analysis comparing genes between stimulated and unstimulated myeloid populations revealed pathways associated with interferon response, inflammatory response, and chemokine migration in stimulated cells (Fig. S10A) suggesting an inflammatory response. Further, key markers of myeloid cell activation and differentiation such as *SLAMF1*, *CD247*, *GBP2*, *CXCL8* and *BATF2* were upregulated after PMA/Ionomycin activation, revealing a woodchuck myeloid-specific cell activation signature (Fig. S11).

**Fig. 4.**
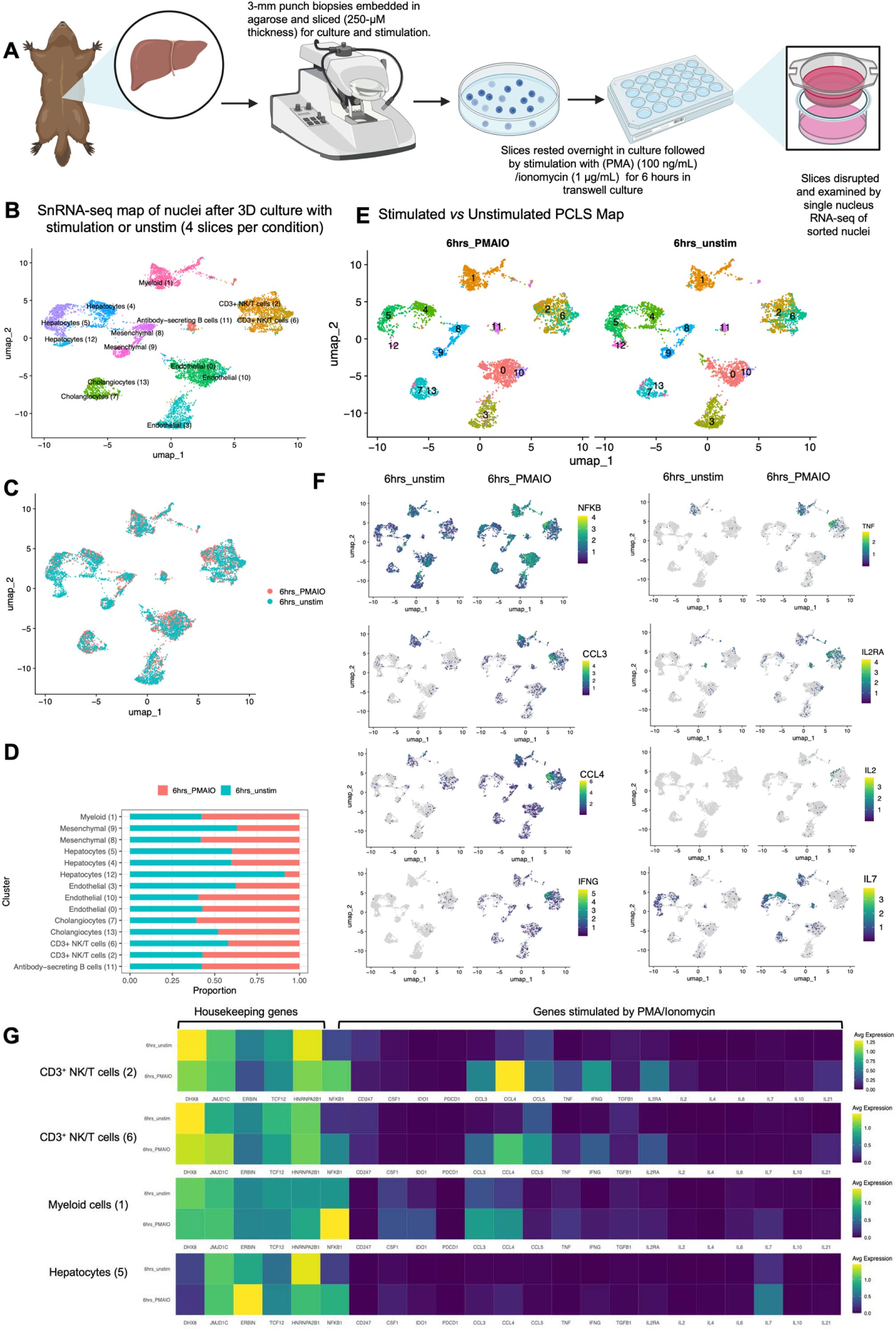
Functional examinations of immune cell populations in 3D Woodchuck liver cultures show upregulated inflammatory signals following PMA/Ionomycin stimulation. (A) Healthy woodchuck precision cut liver slices were stimulated with PMA/ionomycin and single-nucleus RNA sequencing was performed. (B) A UMAP of 9,208 stimulated woodchuck liver cells labelled with their assigned cell identity. (C) A UMAP of stimulated woodchuck liver cells grouped by their sample IDs. (D) A barplot of sample IDs as distributed across clusters labelled in (E) where UMAPs are split by treatment (PMA/ionomycin or PMAIO and unstimulated, respectively). (F) UMAPs of stimulated woodchuck liver cells split by sample ID with heatmaps of key inflammatory genes projected onto the maps. (G) Heatmaps showing the expression of housekeeping genes and genes stimulated by PMA/Ionomycin in four of the snRNA-seq clusters: CD3+ NK/T cells (2), CD3+ NK/T cells (6), myeloid cells (1), and hepatocytes (5).

We extended our characterization of PMA/ionomycin PCLS stimulation to intrahepatic T cells and expected induction of TGFβ signalling and IFNγ activation. Lymphocytes were annotated using the marker genes described above and PMA/ionomycin stimulation resulted in higher expression levels of key T cell activation markers: *NFKB1, CCL3, CCL4, CCL5, TNF,* and *IL2RA* (Fig. 4F,G, Table S10). This response is comparable to the previously reported human PBMC T-cell response to PMA/ionomycin activation, which likewise results in higher levels of *IL2, IFNG, TNF-*α, and *RANTES (CCL5)*.^69,70^ The observed response in woodchucks suggests that their intrahepatic T cells are primed towards TH1 in favour of TH2 or TH17 responses.

Finally, we explored genes activated in B cells, hepatocyte, endothelial cell and mesenchymal cells post-PMA/ionomycin stimulation to identify woodchuck inflammation specific genes in these cell types. All stimulated PCLS cell types display pro-inflammatory NFKB1 pathway activation seen through the upregulation of key cytokines: *CXCL10*, *CCL4* and *CXCL8*, and inflammatory effector proteins *TIFA* and *IDO1* (Fig. 4F,G, Fig. S11, Table S10). Hepatocytes also show increased expression of hepatocyte-inflammation specific genes: *SAA1*, *SAA2*, *B4GALTS*, and *VEPH1* (Table S10).^39^ Furthermore, we see that stimulated woodchuck hepatocytes express *FAS* and cholangiocytes upregulate caspase proteins transcripts *CASP4* and *CASP6* post-stimulation (Table S10), suggesting activation of cell-death programs in liver parenchymal cells as a result of inflammatory signalling. Further, we note an increase in cell adhesion and leukocyte trafficking markers in woodchuck endothelial cells (e.g. *CD44*, *ITGAM*, *SYN3*, and *VCAM1*), pro-survival pathway genes (e.g. *BATF3*, *IRGM*, *TCF3*) and a T cell and endothelial cell specific upregulation of key type I inflammation pathway genes *IFNG* and *STAT3* (Fig 4F,G, Table S10). We also note that cholangiocytes and endothelial cells secrete *CSF1* (Table S10), a monocyte to macrophage differentiation growth factor, to presumably promote macrophage accumulation and activity in inflamed liver tissue. Finally, antibody-secreting B cells and hepatocytes increase expression of lymphocyte growth factor gene *IL7* (Fig 4F,G, Table S10), suggesting a development of a T cell supportive niche in inflammatory liver tissue. Here, we characterise genes and pathways associated with inflammatory signalling in healthy woodchuck tissue to establish key biomarkers of interest for exploration in future liver disease studies.

### Cell level characterization of chronic WHV infection as a pre-clinical model of chronic HBV infection

To expand on the value of the woodchuck as a pre-clinical model of chronic viral liver infection, we applied our above scRNA-seq workflow to cWHV-infected woodchuck livers (Fig 1D, Fig. 5A, Fig. S12A). Using the marker genes identified through the above analysis, 18 populations of parenchymal, stromal and immune cells were annotated when comparing the livers of 2 healthy animals to 3 cWHV animals (Fig. 5B,C, Fig. S12B,C; see diseased animal characteristics in Table S1; see Table S11 for cell-type specific DGE analysis and see Table S12 for a diseased vs healthy DGE analysis for each cell type). Cells from infected livers express WHV viral transcripts while healthy samples do not (Fig. 5D). Both healthy and cWHV samples cluster well without integration, indicating limited biases as result of technical handling of the samples. Interestingly, we identified a cluster of T cells (IFNG_CXCR6_Tcells) with upregulation of key effector (*IFNG, KLRD1, CCL3*) activation (*FUT8*) and exhaustion associated markers (*TOX, NR4A2*) in cWHV that were not present in the healthy controls (Fig. 5E,F). Indeed, relative to the remaining CD3+ T cells, the cWHV-specific IFNG_CXCR6 T cells had both a higher exhaustion and cytotoxic score in both^71^ healthy and cWHV animals (Fig. 5G, Table S13 for gene sets). Furthermore, similar to chronic HBV infection, histological evaluation of cWHV-infected tissue displays immune infiltration, and mild hepatocytic necrosis at the periportal region while the central veins remain undamaged (Fig. S13). These data indicate that CD8+ T cells may be exhausted in cWHV livers leading to impaired viral clearance in chronically infected samples, similarly to recent scRNA-seq mapping of chronic HBV infected human livers.^31^

**Figure 5:**
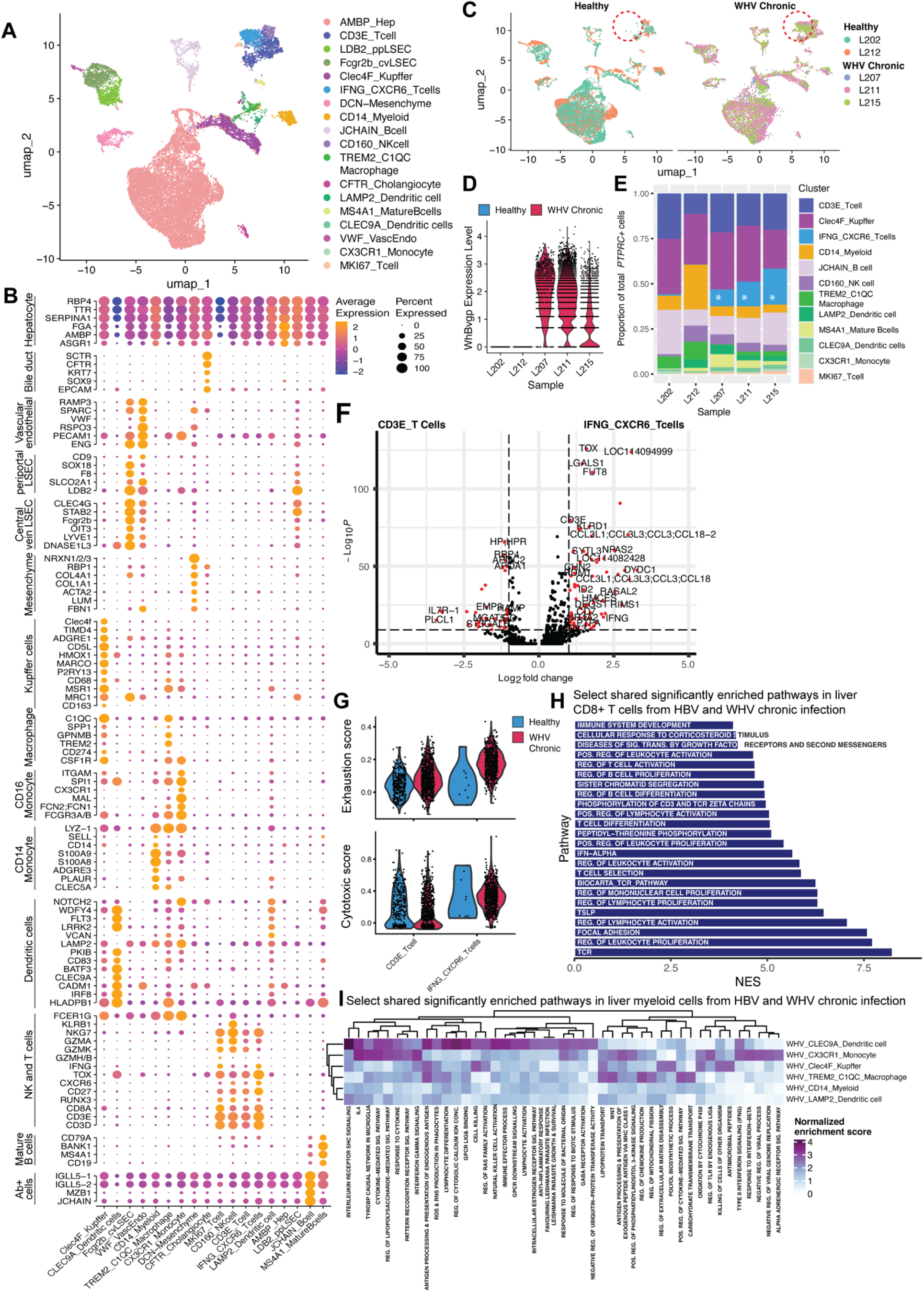
Shared pathways of T cell exhaustion and activation in cWHV-infected woodchuck livers and human hepatic chronic HBV infected tissue. A. Annotated UMAP of clusters generated from healthy (n=2) and diseased woodchuck livers (n=3) resulted in 41 clusters that could be grouped into 18 cell populations. B. Lineage-associated marker genes as described in the previous figures and top-most differentially expressed genes per cluster. C. UMAP split by disease stage and coloured by sample. Subclusters that are enriched in disease vs healthy are highlighted with a red dotted circle. D. Expression of WHV-associated genes is only found in samples from woodchucks chronically infected woodchuck with WHV. E. Stacked bar chart showing the relative proportions of immune cell populations per cluster indicates the emergence of an *IFNG^+^CXCR6^+^ CD8A^+^ T cell* population (indicated with white star) with cWHV infection. F. Volcano plot depicting the differentially expressed genes between the *IFNG^+^ CXCR6^+^ CD3E^+^ CD8A^+^* T cell cluster emerging in WHV infection and all other liver-resident *CD3E^+^*T cells. G. Average expression of cytotoxicity and exhaustion associated gene-sets in T cell populations in healthy and cWHV-infected samples. H. Overlapping enriched pathways in gene-set enrichment analysis between T cells from livers of humans chronically infected with HBV (n=5) (from Zhang et al., *GUT* 2023) and woodchuck livers with cWHV infection (n=3). I. Overlapping enriched pathways in gene-set enrichment analysis between various myeloid populations from livers of humans or woodchucks chronically infected with HBV or WHV.

To determine if cWHV and HBV infections result in similar alterations to immune related pathways in the liver, we sub-clustered T cells from a publicly-available dataset of chronic HBV infected samples in the immune-active phase (GSE182159, Fig. S14A-D).^31^ We then utilized pathway analysis to identify shared pathways enriched in T cells and myeloid cells in both cWHV-infected woodchuck and HBV-infected human tissues (Fig. 5H,I). This analysis showed shared PD-1 activation, lymphocyte activation and pathways associated with myeloid activation such as “IFNG signalling” in chronic WHV carriers -reinforcing the parallels between chronic WHV and chronic HBV. To expand on the PD-1 shared pathway to delineate exhaustion *vs*. activation, we further examined key activation and exhaustion genes in both scenarios and found a similar pattern of upregulated gene expression (*TIGT, TOX, TNFRSF9, CTLA4, IFNG, CXCR6*) (Fig. S15) with the Zhang dataset. Several of these T cell activation genes have also been shown in studies of liver resident T cells in both NASH and in chronic HBV infection.^72,73^ This analysis indicates that cWHV-infected woodchuck livers exhibit disease related pathway activation that have been noted in human chronic HBV infection. This highlights the potential and value of the woodchuck model for testing immunomodulatory interventions to drive antiviral immunity and promoting a functional cure for HBV.

## Discussion

We have generated the first atlas of woodchuck parenchymal and non-parenchymal liver cells and PBMCs using single-cell RNA-sequencing. Having such a resource enhances the translational potential of this unique, clinically valuable animal model that has previously been limited by a lack of molecular biology tools. This atlas will allow for the examination of parallels between WHV and human HBV pathogenesis as a stepping stone to generate new therapeutics.

In our analysis, hepatocyte zonation could be largely identified in the single-cell dataset due to the clear presence of pericentral markers determined from the spatial transcriptomics data. This data type was crucial for determining pericentral and periportal hepatocyte markers that were much more distinct in the spatial transcriptomics compared to the scRNA-seq. In the scRNA-seq data, ambient RNA was largely composed of periportal hepatocyte marker genes making these cells more difficult to identify. The additional cluster of “Unknown Hepatocytes” appears to be unrelated to zonation and may be interesting to explore further.

Broad similarities were observed between human and woodchuck cells across most other clusters identified (Fig. S3A-D). We were able to successfully use lineage specific markers (monocytes/macrophages (*CD14/CD68*), T cells (CD3 isoforms), B cells (*CD19/MS4A1*)) from mouse and human to help identify the majority of the immune clusters from woodchuck liver and blood. Furthermore, many of these clusters were also transcriptionally similar to immune cell subsets commonly observed in humans. For example, our analysis of woodchuck PBMCs revealed three monocyte subclusters based on the differential expression of *CD14* and *FCGR3A (CD16)*. These subpopulations had similar gene expression profiles to different populations of human monocytes^74^, suggesting translatable immune targets between woodchucks and humans.

We also identified several myeloid populations including more monocytic and more tissue-resident cells in the woodchuck liver which mirrored the transcriptional landscape previously reported in humans.^5,26^ The similarities in monocyte/macrophage populations between human and woodchuck may be of particular importance when considering potential future treatment strategies targeting these populations. Indeed, tumour associated macrophages represent a promising therapeutic target given their important role in promoting tumour development, progression, and metastasis in many cancers.^75–77^ Overall, the parallels observed between human and woodchuck cells further support the potential of the woodchuck as a model for HCC development from chronic HBV infection in humans.

In annotating our atlas, we emphasise the conservation of marker genes and expected cell populations from previous single-cell analyses of the human liver. Although these orthologous relationships allowed us to describe the woodchuck liver and blood in detail, it is important to note that most of our conclusions are derived from the assumption that similar peptide sequences across translated genes are functionally similar. Functional validation tests, along the lines of the precision cut liver slice stimulations, can provide evidence of activation of gene pathways in intrahepatic woodchuck cells we have described.

A key challenge related to working with total liver homogenates is the fragility of hepatocytes which leads to high cell death and consequent release of ambient RNA, which can complicate annotation. Filtering the liver homogenate datasets with DropletQC greatly improved the clarity of signals from the liver tissue data. Liver samples generally contain high levels of ambient RNA that connect clusters with a gradient of ambient RNA expression. These “gradient clusters” are difficult to interpret and may be falsely interpreted as interacting cells or differentiation gradients. DropletQC identified many of these “gradient clusters” as empty droplets due to their low nuclear fraction. It also removed clusters that were previously thought to be periportal hepatocytes, but also suffered from large amounts of ambient RNA and batch effects.

Our single-cell examination of healthy and cWHV-infected woodchuck livers identified shared cellular programs between the woodchuck, human, and murine liver cellular landscapes and found exhaustion-like antiviral immunity parallel to human HBV infection. However, the WHV-infected woodchuck model of HBV is not without its caveats. Chronic HBV infection in humans is frequently associated with progressive liver fibrosis or cirrhosis; however, WHV-infection in woodchucks very rarely causes liver fibrosis and cirrhosis does not develop.^14,78,79^ Moreover, progression to cWHV-induced HCC is more consistent and present in 90% of animals with chronic hepatitis within 18 to 36 months after infection with WHV^14,80^, compared to humans where approximately 5% of chronically HBV infected individuals develop HCC^81^. Finally, WHV DNA randomly integrates to the liver genome, although insertions around the *N-myc2* locus were also found, in woodchuck HCC^80,82,83^, whereas HBV insertional mutagenesis is largely thought to be random in humans.^84^ Thus, findings related to WHV-triggered HCC development in woodchucks may not fully replicate processes expected to occur in human HBV-induced liver cancer. While previous bulk-analyses have identified intrahepatic expression of markers associated with T cell exhaustion and inhibition of cytokine signalling in livers of woodchucks with cWHV hepatitis^85^, here we describe the transcriptome of a cWHV-infected liver derived exhausted-like T cell population associated with viral persistence. Importantly, our data suggests shared dynamics of T cell response in cWHV and HBV thereby reinforcing the value of this model for immune modulating studies to induce potent antiviral immunity facilitating sterilizing HBV clearance.

Our work has characterised the woodchuck model with single cell resolution through transcriptional mapping of liver and peripheral blood mononuclear cells. Specifically, the high-resolution characterisation of immune cells allows for hypothesis generation of potential immunotherapies that can then be tested and analysed in the woodchuck-WHV infection model. Future directions of this work may include adding samples from other stages of WHV infection and forms of WHV hepatitis,, and incorporating more data types at the single-cell level (e.g., ATAC-seq) to gain a deeper understanding of cell characteristics and their contribution to disease progression or resolution. In addition, certain clusters will benefit from validation experiments to confirm the cellular phenotypes we predicted in these maps. Taken together, these maps will enhance the utility of the woodchuck-WHV model and benefit the HBV community by acting as a reference for examining the liver cellular microenvironment and intrahepatic immune responsiveness to test antiviral and immunotherapies through the course of WHV-induced inflammatory liver disease and cancer.

## Abbreviations

scRNA-seq: single-cell RNA sequencing
snRNA-seq: single-nucleus RNA sequencing
HCC: hepatocellular carcinoma
ATAC-seq: assay for transposase-accessible chromatin using sequencing
PBMC: peripheral blood mononuclear cell
HBV: hepatitis B virus
WHV: woodchuck hepatitis virus
HSC: hepatic stellate cell
LSEC: liver sinusoidal endothelial cell
PCLS: precision cut liver slice
DE: differentially expressed
GSEA: Gene Set Enrichment Analysis
TLH: tissue liver homogenate

## Data Availability

The woodchuck genome sequence and annotation are available at https://doi.org/10.5281/zenodo.10855128. The raw RNA-seq data and data matrices output by Cell Ranger are available at the Gene Expression Omnibus (GEO) at accession numbers GSE264104 (scRNA-seq), GSE264107 (spatial), and GSE264112 (snRNA-seq). The single-cell and single-nucleus datasets are hosted by UCSC Cell Browser at https://woodchuck-liver.cells.ucsc.edu. The spatial transcriptomics data are available for exploration at https://macparlandlab.shinyapps.io/woodchuck-spatial-v2/.

The code used to process the data can be found at https://github.com/14zac2/HealthyWoodchuckMap.

## Acknowledgements

The authors acknowledge the Princess Margaret Genomics Centre, the Pathology Research Program and the Advanced Optical Microscopy Facility at University Health Network for their support and services. The graphical abstract and select figures were created with Biorender.com.

