## Supplementary Materials for "A Single-Cell Woodchuck Liver Atlas Identifies Healthy and Disease-related Cellular Programs Conserved in Human"

**A single-cell atlas of the woodchuck liver maps an immune landscape and hepatocyte zonation patterns reflective of the human liver**

Zoe A. Clarke, Xinle Wang, Jawairia Atif, Lewis Y. Liu, Lawrence Wood, Damra Camat, Sai Chung, Xue-Zhong Ma, Justin Manuel, Si Lok, Timothy Lau, Cornelia Thoeni, Tomasz I. Michalak, Ian D. McGilvray, Gary D. Bader, Sonya A. MacParland

### Supplementary methods

#### Woodchuck liver tissue perfusion

Animals were sedated by intramuscular injection of mixture of 20 mg/kg ketamine hydrochloride and 5 mg/kg xylazine, then inhalant anaesthesia with 2-4% isofluorane. A single intramuscular injection of butorphanol tartrate at 0.3 mg/kg was then administered. Tissue was dissociated with warm 1X Hanks' balanced salt solution containing collagenase MA (VitaCyte) and neutral protease (VitaCyte). 4 mL of dissociation solution was added to tissue fragments and the tissue was incubated at 37°C for 10 minutes. Cell suspensions were then homogenised using the gentleMACS tissue dissociator and filtered through a 70-µm filter while on ice. 5% of fetal bovine serum (FBS) and PBS at 1:1 ratio were added into the cell suspensions for enzyme inactivation.

#### Woodchuck Precision Cut Liver Slice culture

5% low-temperature agarose gel was prepared by dissolving 1 g of agarose (Sigma-Aldrich- A0701) in 10 mL of MilliQ water followed by the addition of 10 mL of pre-warmed Williams E Media. Biopsies were placed in peelable base molds and overlaid by the liquid agarose gel without displacing the tissue. After solidification, base molds were peeled to free the agarose blocks. Excess agarose on the sides of the block was trimmed and tissue was glued onto the cutting platform with surgical glue (3M Vetbond, 1469Sb).

#### Single-nucleus RNA-sequencing

A 2X stock solution of salt-Tris solution (ST buffer) was made with 292 mM NaCl (Sigma-Aldrich), 2 mM CaCl2 (Sigma-Aldrich), 42 mM MgCl2 (Sigma-Aldrich), and 20 mM Tris-HCl, pH 7.5 (Thermo Fisher Scientific) in ultrapure water (Thermofisher).

1X ST buffer was prepared via an additional 1:1 dilution with Ultrapure water. The CST detergent buffer containing 50% 2X ST stock solution, 0.0049% CHAPS (Sigma-Aldrich), 0.0001% BSA (New England BioLabs) was used for isolation. Each sample was diced by spring scissors for 10 minutes in the CST buffer. The solution was washed with 1X ST through a 40-µM mesh filter (Falcon), and spun down at 500 x g.

#### Mapping and annotating a reference Woodchuck whole genome

The woodchuck genome was sequenced with a combination of 77X long-read Pacbio and 120X short-read Illumina sequencing from woodchuck sample L192, sourced from Labrador, Canada, and acquired and preserved by the Molecular Virology and Hepatology Research Group, Faculty of Medicine, Memorial University of Newfoundland, St. John's, NL, Canada. Our genome is 2,638,837,009 bp and is composed of 3,381 contigs assembled into 2,683 scaffolds, with a contig N50 of 10,671,126 and a scaffold N50 of 82,176,798. Previous woodchuck genome assemblies by Puiu *et al.* (2020)[1] and Alioto *et al.* (2019)[2] have 8,860 contigs (no scaffolds) with a contig N50 of 1,094,236, and 48,534 scaffolds with a scaffold N50 of 892,188, respectively, indicating that our assembly is much more complete. Since beginning this project, an earlier version of this genome has been published on Refseq and is the current woodchuck reference genome (assembly GCF_021218885.2). Following this, a very similar version of the woodchuck genome used for this project was uploaded to Genbank as assembly GCA_024703135.1.

The woodchuck genome used for this project was annotated using a combination of gene annotation liftover from related genomes, RNA-seq, and a novel in-house exon homology-based gene prediction program (unpublished) (Sup. Fig. 11). Gene annotation liftover was performed with Liftoff[3], a tool that aligns gene sequences across genomes regardless of the structural differences between them at the contig/scaffold level. We lifted annotation from RefSeq annotation sources of three closely related species, the yellow-bellied marmot *(Marmota flaviventris)*, Alpine marmot *(Marmota marmota)*, and the 13-lined ground squirrel *(Ictidomys tridecemlineatus)*. These species were chosen because they were phylogenetically close to the woodchuck and had genomes available on NCBI's RefSeq database, with respective RefSeq version IDs: GSC_YBM_2.0, marMar2.1, and SpeTri2.0. Liftoff was run under default parameters with the additional parameters "-flank 0.5 -copies" to align 50% of gene flanking sequences and look for additional orthologs in the woodchuck genome from the original genes in the reference species.

Woodchuck RNA-seq reads from seven different tissues (liver, kidney, spleen, lung, heart, pancreas, and thymus with respective NCBI Sequence Read Archive accession codes SRR10172922, SRR10172923, SRR10172924, SRR10172925, SRR10172926, SRR10172930, and SRR10172931)[2] were each separately aligned to the woodchuck genome with HISAT2 v2.2.1[4] and used to predict transcript location and structure with StringTie v2.1.3[5]. HISAT2 was run with the "--dta" flag to tailor the alignments for transcript assemblers such as StringTie, and the resulting SAM files were converted to sorted BAM files with samtools[6] ("samtools view" with parameters "-Shu" followed by "samtools sort"). StringTie was run with default parameters. The seven resulting annotation sets were combined and filtered to create a unified genome annotation using the Mikado v2.0rc2 (with Python v3.6.10) pipeline[7], an automated tool designed to mimic manual gene curation from multiple transcript assemblies. Mikado drew upon evidence from the following additional tools: open reading frames (ORFs) were determined with Transdecoder v5.5.0[8] (using commands "TransDecoder.LongOrfs" with the argument "-m 30", and "TransDecoder.Predict" with the argument "--retain_long_orfs_length 30" as recommended in the Mikado documentation); splice site validation was performed with Portcullis v1.2.0[9] (default parameters); and candidate transcripts were compared to the Swiss-Prot database from UniProt[10] (accessed Dec. 24, 2020) with BLAST+ v2.11.0[11] to estimate homology ("blastx" with parameters "-max_target_seqs 5 -outfmt "6 qseqid sseqid pident length mismatch gapopen qstart qend sstart send evalue bitscore ppos btop" -evalue 0.000001").

Mikado relies on tailored configuration and scoring files which input rules on how the program should be filtering and prioritising the different possible transcripts. To filter the seven RNA-sequencing-based annotations produced by StringTie, the configuration and scoring files were customised following Mikado guidelines on how to filter noisy RNA-seq data (mikado_round1_configurationFile.yaml, mikado_round1_scoringFile.yaml, customised following guidelines from: https://mikado.readthedocs.io/en/stable/Tutorial/Adapting/). Specifically, we created a configuration file with the following rules: introns which are 60 - 70,000 base pairs are favoured (based on patterns seen in similar species); the flanking region of a transcript is increased to 2,000 base pairs to identify and filter out more fragments; chimeric transcripts that are not supported by a good BLAST hit are split; the minimum score for calling an alternative splicing event is increased to 0.75; transcripts with retained introns are not counted as valid alternative splicing events; and there is a maximum of five isoforms per transcript.  The scoring file was modified from mammalian.yaml, which was the default mammalian scoring file that was included with this version of Mikado. Modifications included: discarding any multiexonic transcripts without introns verified by Portcullis; discarding splicing events that would be canonical on the opposite strand; requiring a transcript to have a coding region of at least 300 base pairs; discarding alternative splicing events that have more than one open reading frame; and considering open reading frames in coding sequences as fragments if they are less than 300 base pairs long. Intron lengths were restricted to 30 - 900,000 base pairs and the possible lengths of 3' and 5' untranslated regions were increased to 9,000 and 6,000 base pairs respectively, to reflect the distribution of lengths seen in the genomes of closely related species. Transcripts that passed all filters and were deemed most likely to be real based on the configuration and scoring files were kept as RNA-sequencing evidence for our gene annotations.

A partial set of conserved genes (WCK01_AAH20201022_F8-SC2.anno20210519.gff3.gz) was also derived from an in-house exon homology-based gene prediction program (Lok, Lau, unpublished). This annotation set was combined with the Liftoff genome annotations from our three selected closely related species and the RNA-seq-based annotation from StringTie filtered using a second application of the Mikado pipeline. We used different settings for this Mikado run, relaxing some filtering rules that were applied in the first round when processing highly fragmented raw RNA-seq data (mikado_round2_configurationFile.yaml, mikado_round2_scoringFile.yaml). For instance, we did not rely on splice site validation by Portcullis, although the parameters for Transdecoder and BLAST+ were identical to the first round of Mikado filtering.

In addition to removing splice site validation by Portcullis, the configuration and scoring files were customised for the second round of Mikado. The configuration file was modified from the default provided by Mikado, by again increasing the flanking region of transcripts to identify gene fragments to 2,000 base pairs; chimeric transcripts with ORFs matching two different BLAST hits were split, the minimum score for an alternative splicing event was increased to 0.99; confirmed introns were not required; retained introns were now valid alternative splicing events; and each transcript could have a maximum of five isoforms. The input annotation sets were also ranked to prioritise transcripts from higher quality sources. The in-house homology-based gene annotation was given a score of ten and considered as a reference annotation, the gene-liftover-derived annotations were given a score of five, and the filtered, RNA-seq annotation was given a score of three. The scoring file was modified again from mammalian.yaml provided by Mikado: minimum intron length was increased to 30 base pairs; the 3' and 5' untranslated regions were increased to support a maximum of 9,000 and 6,000 base pairs respectively; alternatively spliced transcripts must have a maximum of one open reading frame; and transcripts in flanking regions must have a coding region of over 300 base pairs to not be considered a fragment.

Genome annotation quality and accuracy was assessed by comparing fourteen genome annotation evidence sources with GFFCompare v0.11.2[12] (specifically results from StringTie, Augustus, Liftoff, and the first round of Mikado) and analysing gene and transcript structure with Mikado. Our in-house gene set, which consists of a partial annotation set of highly accurate genes, was used as a reference against which the other annotation sets were compared to estimate the accuracy of each annotation set. GFFCompare measures the number of identical base pairs, exons, introns, and full transcripts that occur between multiple annotation sets of the same genome. Gene structure was calculated using Mikado's "util stats" command, and indicates features such as exon length, exons per transcript, and splice variants per gene. The woodchuck summary statistics were compared to the summary statistics of the yellow-bellied marmot, Alpine marmot, and 13-lined ground squirrel, to ensure that the filtering by Mikado selected transcripts with a distribution of features similar to those from closely related species.

The resulting set of protein-coding genes was translated with GFFRead v0.12.1[12] (parameters "-y -S"). Translated reads were compared with the peptide sequences from human (RefSeq version GRCh38), mouse (RefSeq version GRCm39), yellow-bellied marmot, Alpine marmot, and 13-lined ground squirrel for protein orthology detection. This was done with OrthoFinder v2.5.1[13], a tool that infers orthology using a species phylogeny. We provided OrthoFinder with a known species phylogeny derived from phyloT v2[14], an online resource that constructs phylogenetic trees based on NCBI taxonomy, to improve the accuracy of the orthology predictions[13]. Resulting pairwise orthology was used to pair the translated woodchuck genes with the proteins from the other input species. These protein-protein relationships were converted to gene-gene relationships by associating the translated gene/protein sequences to the encoding gene; this information was taken from the annotation and sequence files associated with the particular version of each genome. One-to-many relationships were indicated by separating human genes with a semicolon where human paralogs exist for a single woodchuck gene. Many-to-one relationships were also incorporated by identifying multiple woodchuck genes to the same human gene where woodchuck paralogs exist. Both methods were combined to accommodate many-to-many gene relationships.

Woodchuck genes were labelled with their corresponding orthologs to support further analyses, with priority given to human orthologs. If a human ortholog did not exist, the mouse ortholog would be used, or yellow-bellied marmot, Alpine marmot, and 13-lined ground squirrel in priority order. Duplicate gene symbols for many-to-one relationships were appended with a suffix (e.g. -1) to maintain the symbol as a unique identifier. The gene name "Btbd35f14;Btbd35f16;Btbd35f18;Btbd35f27;Btbd35f3;Btbd35f4;Btbd35f26;Btbd35f19;Btbd35f20;Btbd35f17;Btbd35f9;Btbd35f11;Btbd35f10;Btbd35f15;Btbd35f13;Btbd35f12;Btbd35f25;Btbd35f5;Btbd35f21;Btbd35f23;Btbd35f29;Btbd35f28;Btbd35f22;Btbd35f7;Btbd35f6;Btbd35f8;LOC118568425;Btbd35f1;Btbd35f2" was shorted to "LOC118568425", as it represented a one-to-many relationship with the "many" genes exceeding the maximum gene name character count of 256 by CellRanger.

### Supplementary results

#### Mapping and annotating a reference Woodchuck whole genome

At the start of our project, no woodchuck genome existed against which to map our genomics data. Mapping to the closest well-annotated genome, alpine marmot *(Marmota marmota)*, resulted in poor quality scRNA-seq cell annotation (approximately 50% aligned transcripts). We thus sequenced the woodchuck nuclear and mitochondrial genomes using a PacBio Sequel DNA long read sequencer (77 times genome coverage) for primary assembly, followed by correction of errors with 120x Illumina HiSeq X DNA sequencing coverage (The Centre for Applied Genomics, Toronto). Assembly resulted in a 2.7Gb genome with 2685 scaffolds, and a scaffold N50 of >82Mb (a measure of mean scaffold length, weighted towards longer scaffolds), deposited in GenBank/NCBI Genome as Bioproject number PRJNA797297. We performed manual and automated gene annotation based on woodchuck RNA-seq data from seven tissues (liver, kidney, spleen, lung, heart, pancreas, and thymus)[2] and mapping annotation from closely related species (yellow-bellied marmot, alpine marmot, thirteen-lined ground squirrel) using Liftoff[3] to define 26,645 protein-coding genes and 632 non-coding genes (62,914 mRNA and 705 ncRNA transcripts, considering alternative splicing), including 21,131 orthologs to related species. The woodchuck mitochondrial genome also had 13 protein-coding genes and one non-coding gene annotated. 18,578 woodchuck genes on the nuclear genome and the 13 protein-coding mitochondrial genes were assigned a human ortholog, allowing these genes to be used for pathway analysis derived from the relationships between human genes. There were also 15,846 one-to-one genes found between woodchuck and human, and 15,752 between woodchuck and mouse, which were used for the direct comparison of gene expression across these organisms (Table S14). There were 2021 human gene symbols and 4,313 mouse gene symbols that were not detected in the woodchuck genome (Table S15). These genes were likely either unannotated in annotated genomes we transferred, too divergent to be recognized as an ortholog, or do not exist in the woodchuck. Mapping our woodchuck scRNA-seq data to this genome resulted in roughly 15% more reads aligned to the transcriptome and 7% more genes identified, compared to mapping to the alpine marmot genome.

### Supplementary figures

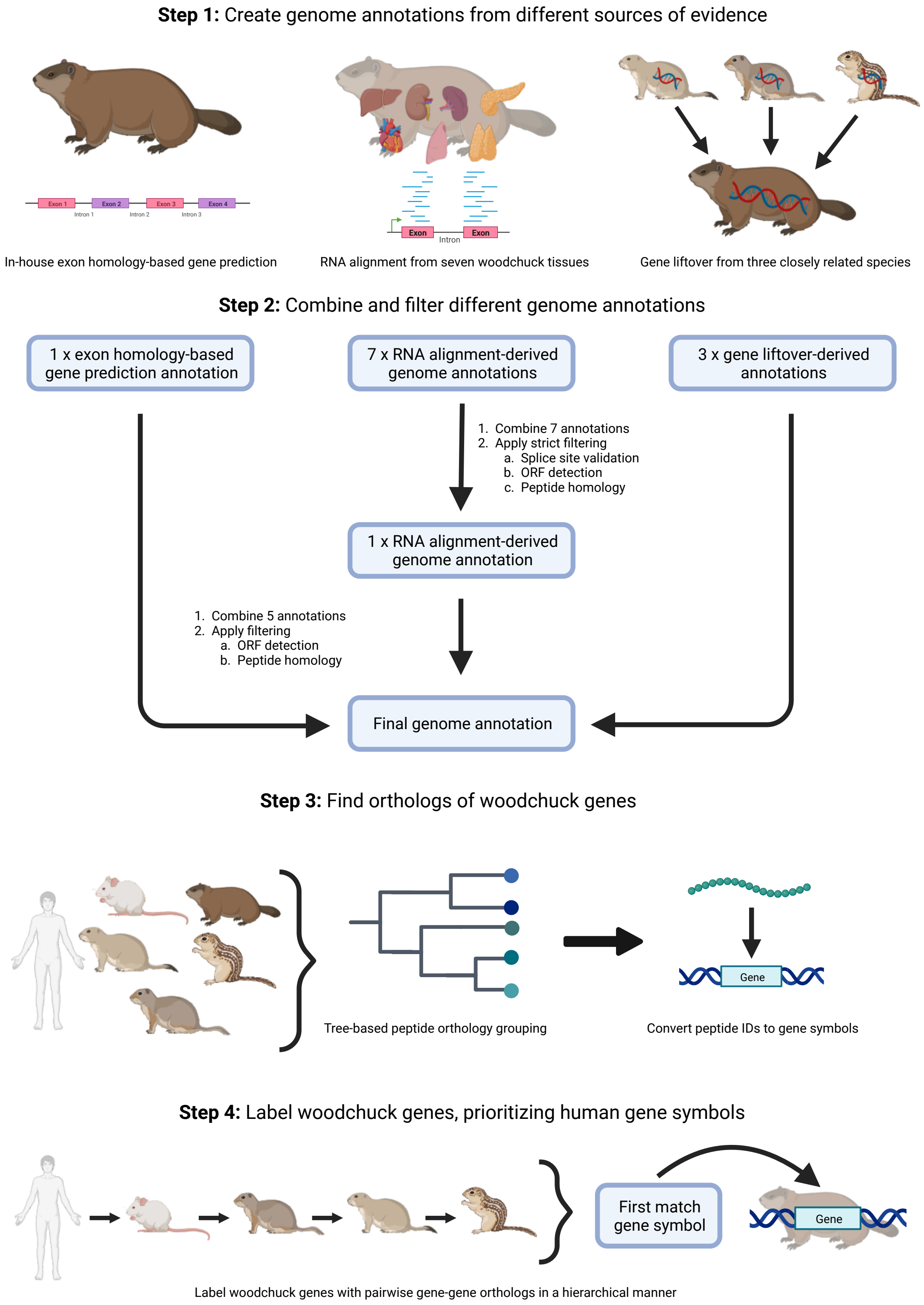

**Supplementary figure 1: Different sources of evidence were used to annotate the woodchuck genome.** Different sources of evidence were used to annotate the woodchuck genome, the process of which is summarised in this figure. The first step was to create separate genome annotations from three different methods: an in-house exon homology-based gene prediction program; RNA alignment using HISAT2[4] and genome annotation prediction with StringTie[5]; and gene liftover using Liftoff[3]. The RNA alignment and genome annotation prediction was performed using RNA from seven different tissues: liver, kidney, lung, pancreas, heart, spleen, and thymus. This created seven separate genome annotation files. Gene liftover was performed on Refseq genomes from three closely related species, creating three genome annotation files: the yellow-bellied marmot *(Marmota flaviventris)*, Alpine marmot *(Marmota marmota)*, and the 13-lined ground squirrel *(Ictidomys tridecemlineatus)*. The second step was to combine and filter all of the different genome annotations using the tool Mikado[7]. Within this step, the seven RNA alignment-derived genome annotations were subjected to strict filtering due to the fragmented nature of the annotation, including splice site validation[9], ORF detection[8], and peptide homology analysis[11]. This created one filtered RNA alignment-derived annotation which was subsequently combined and filtered with the other four genome annotations. The filtering parameters within Mikado were less strict as the annotation sets were less fragmented, and only ORF detection and peptide homology analysis were performed. This created a final genome annotation of 26,645 protein-coding genes and 632 non-coding genes. For the third step, peptide sequences from human, mouse, and the same three closely related species were fed into OrthoFinder[13] which grouped peptides together based on orthology. Peptide-peptide relationships were then converted to gene-gene relationships. Finally, step four involved labelling the genes in the woodchuck genome with the pairwise gene-gene orthologs in a hierarchical manner, prioritising human gene symbols. The final genome annotation was labelled with 21,131 orthologous gene symbols.

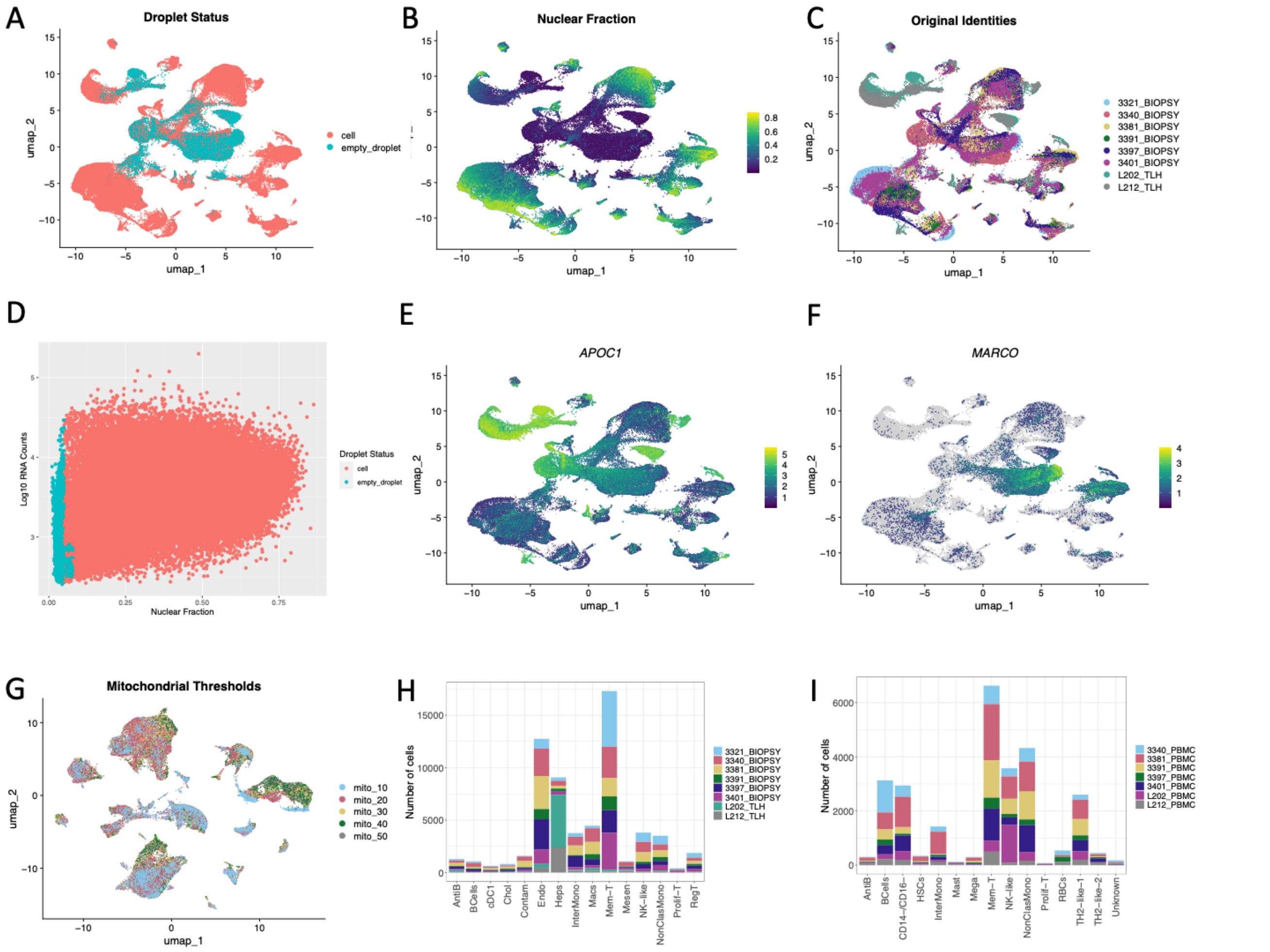

**Supplementary figure 2: Quality control metrics used to filter cells from the woodchuck liver.** QC metrics used to filter cells from the woodchuck liver. (A) Cell (n = 71,962) and empty droplet (n = 28,633) labels provided to woodchuck liver cells from DropletQC under its default settings. The cells are presented on a UMAP of samples merged after the mitochondrial filter was applied, but before DropletQC was applied. DropletQC was used on each sample before merging, and droplets were removed before the final map was integrated. (B) The nuclear fraction of each cell presented on the merged, pre-DropletQC-filtered UMAP of woodchuck liver cells. The nuclear fraction represents the proportion of introns in a particular cell. (C) The original identities of the cells before filtering the sample with DropletQC. (D) A scatterplot comparing the nuclear fraction to the log 10 of RNA counts of each cell. Cells with low RNA counts and low nuclear fractions are considered empty droplets (blue) and filtered out of the sample. (E) *APOC1* expression projected as a heatmap on a UMAP of merged woodchuck liver cells. High levels of *APOC1* represent hepatocytes or contaminated cells which tend to be filtered out of the sample. (F) *MARCO* expression projected onto the merged woodchuck liver cells, roughly indicating the presence of tissue-resident macrophages. Tissue-resident macrophages tend to exhibit higher levels of hepatocyte contamination and are often labelled as empty droplets, and filtered out of the sample. (G) The filtered, integrated woodchuck liver map categorized by different mitochondrial content levels. The number following “mito_” is the maximum percentage of mitochondrial DNA found in a cell. Mitochondrial content below 50% was acceptable, with cells below this threshold being included in the final map. (H) Woodchuck liver cells from Fig. 2A grouped by sample identity in a bar plot. (I) Woodchuck PBMCs from Fig. 2C grouped by sample identity in a bar plot.

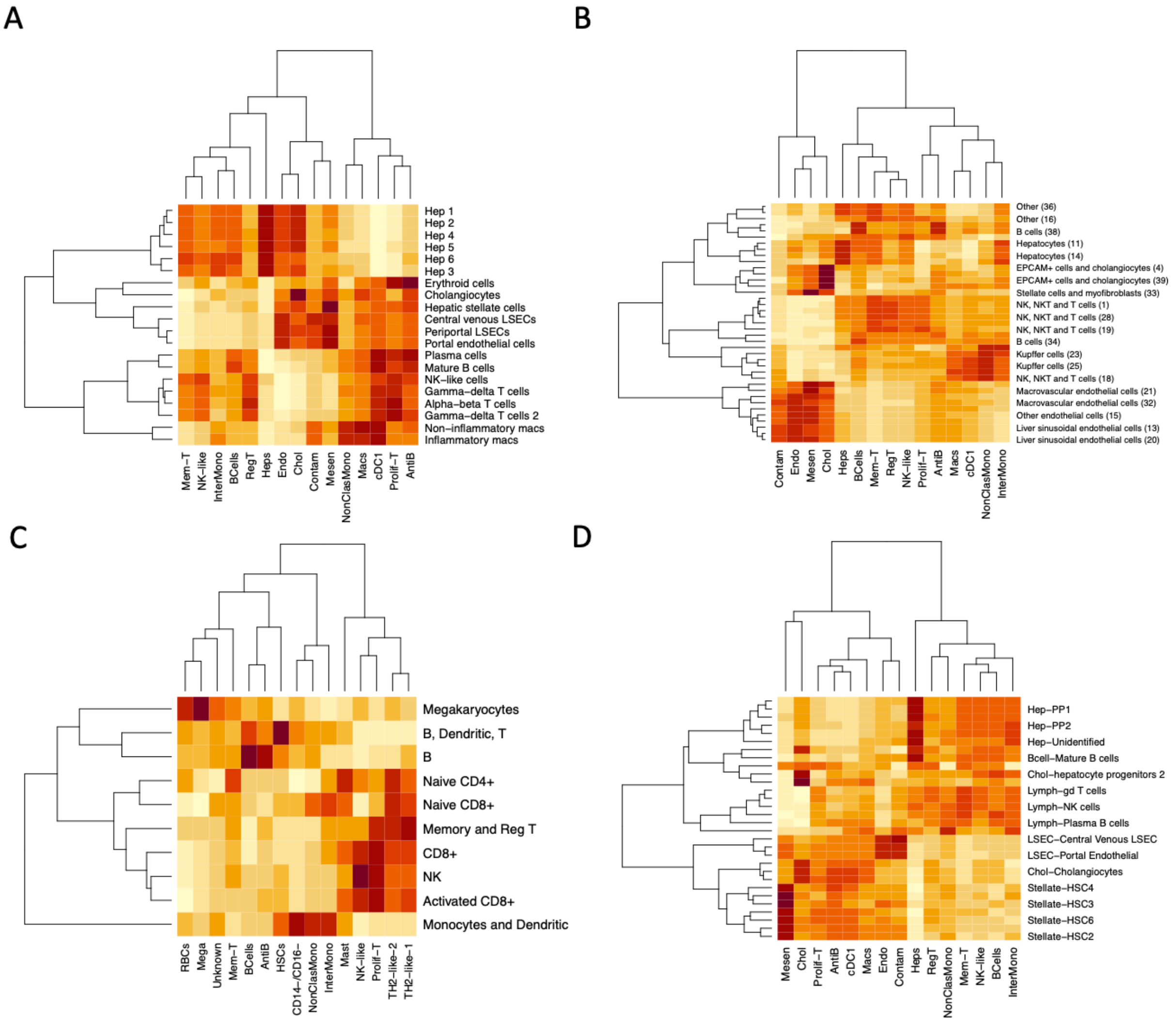

**Supplementary figure 3: Correlation tests demonstrate similarities between cell clusters.** (A) Spearman correlation of woodchuck liver data (x-axis) with the human liver dataset from MacParland et al. 2018[15] (y-axis). This correlation used the 560 of 2,000 genes from the "scale.data" slot in woodchuck that were present as one-to-one orthologs in the human data. (B) Spearman correlation of woodchuck liver data (x-axis) with the human liver dataset from Aizarani et al. 2019[16] (y-axis). This correlation used 628 of the 2,000 genes from the "scale.data" slot in woodchuck. (C) Spearman correlation of woodchuck PBMCs (x-axis) with the 68k PBMC dataset from 10X Genomics[17] (y-axis). This correlation used 165 of the 2000 genes from the "scale.data" slot. (D) Spearman correlation of woodchuck liver data (x-axis) with the human single-nucleus RNA-seq dataset from Andrews et al. 2022[18] (y-axis). This correlation used 630 of the 2,000 genes from the "scale.data".

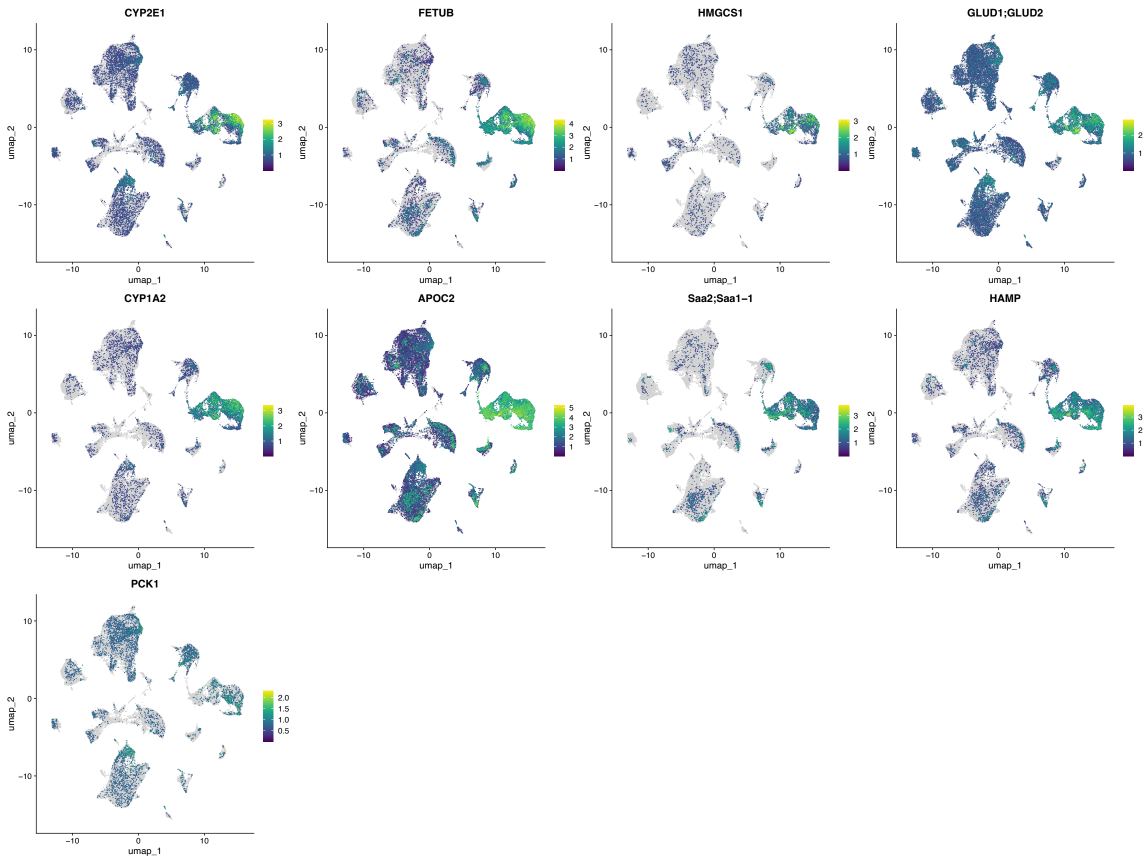

**Supplementary figure 4: Hepatocyte markers from spatial transcriptomics and the single-cell liver map.** Pericentral hepatocyte markers from the spatial transcriptomics data include *CYP2E1, FETUB, HMGCS1, GLUD1;GLUD2,* and *CYP1A2.* Top periportal markers are *APOC2, Saa1/Saa2,* and *HAMP. PCK1* is a periportal marker gene that contrastingly is not highly expressed in the spatial transcriptomics data, but is a clearer indication of periportal hepatocytes in single-cell data.

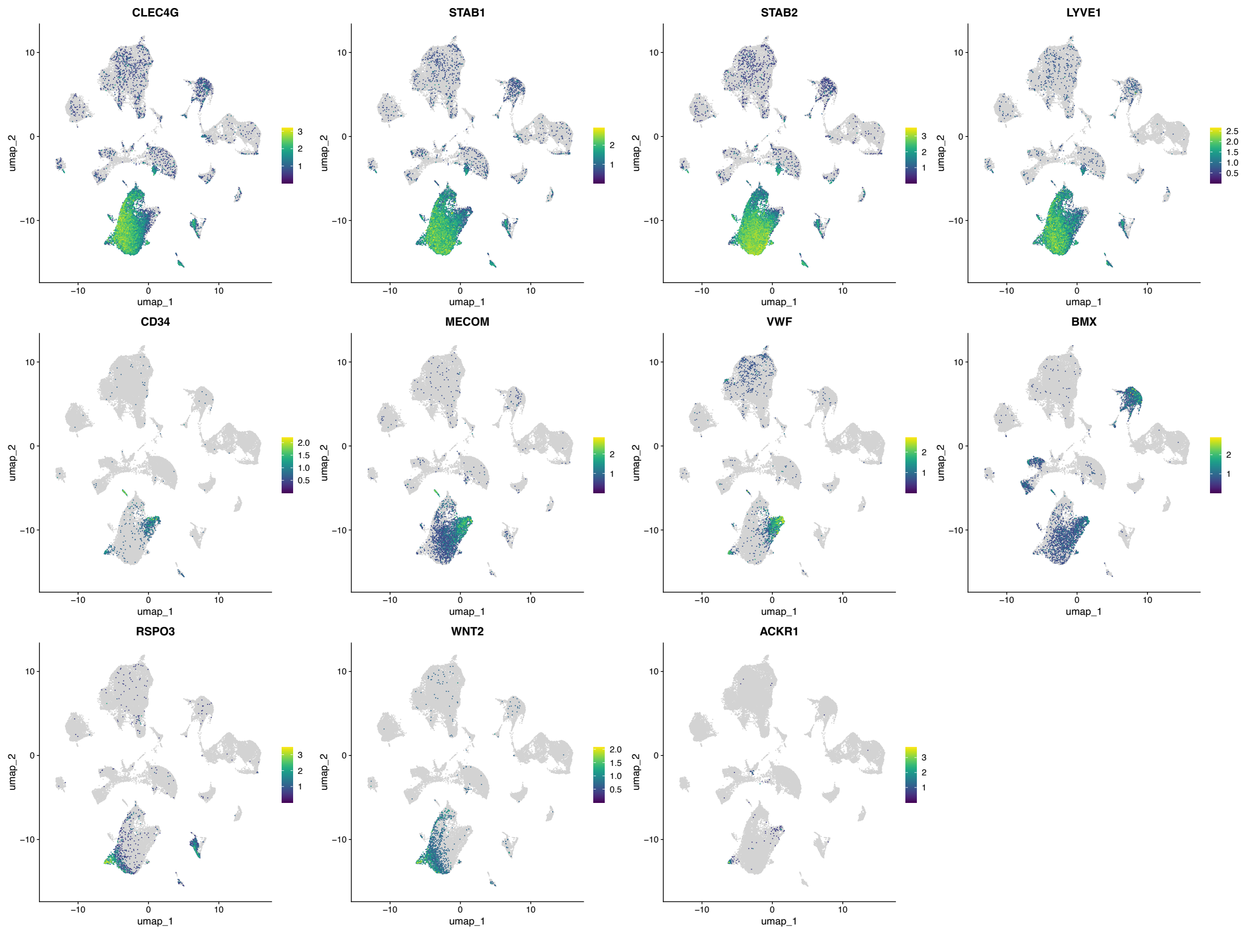

**Supplementary figure 5. Known endothelial cell markers distinguish woodchuck endothelial cell populations.** Significant endothelial cell genes with their expression values plotted onto the single-cell woodchuck liver map. Genes distinguishing the general endothelial cell population include: *CLEC4G, STAB1, STAB2*, and *LYVE1.* A subpopulation of hepatic artery endothelial cells can be distinguished with: C*D34, MECOM, VWF,* and *BMX.* An additional subset of central venous endothelial cells can be distinguished with *RSPO3, WNT2,* and *ACKR1.*

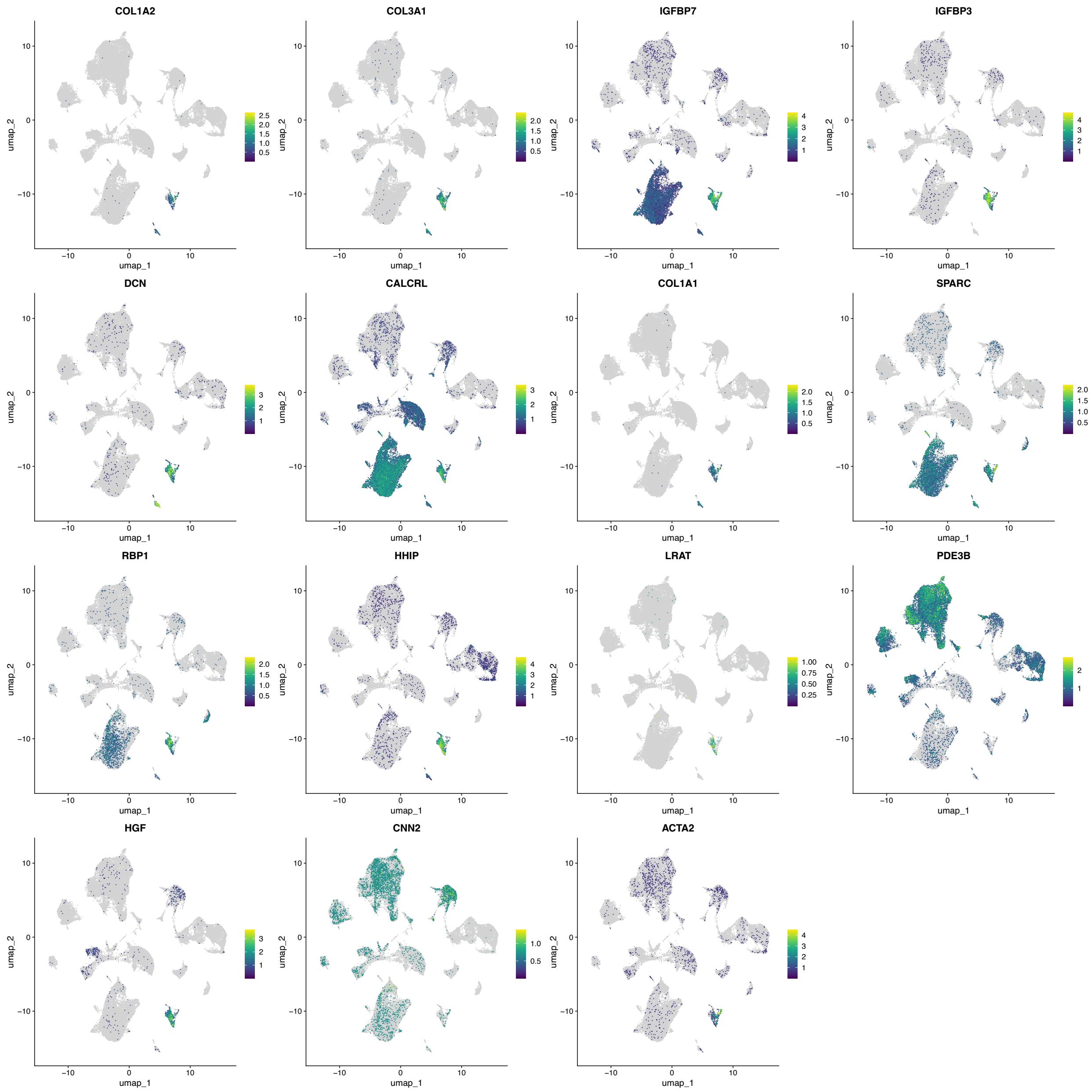

**Supplementary figure 6. Mesenchymal cell markers delineated by scRNA-seq are shared between humans and woodchucks.** UMAP projections of differentially expressed genes from different cell populations in the woodchuck liver. Gene expression values were projected onto the UMAP as a heatmap. Differentially expressed genes in the mesenchymal cell population in the healthy woodchuck liver. All genes were previously found expressed in human mesenchymal cells except *HHIP,* which is a very strong marker in woodchucks. Quiescent mesenchymal cell markers in humans include *LRAT* and *PDE3B*, and activated mesenchymal cell markers include *ACTA2, SPARC*, and *COL1A1.*

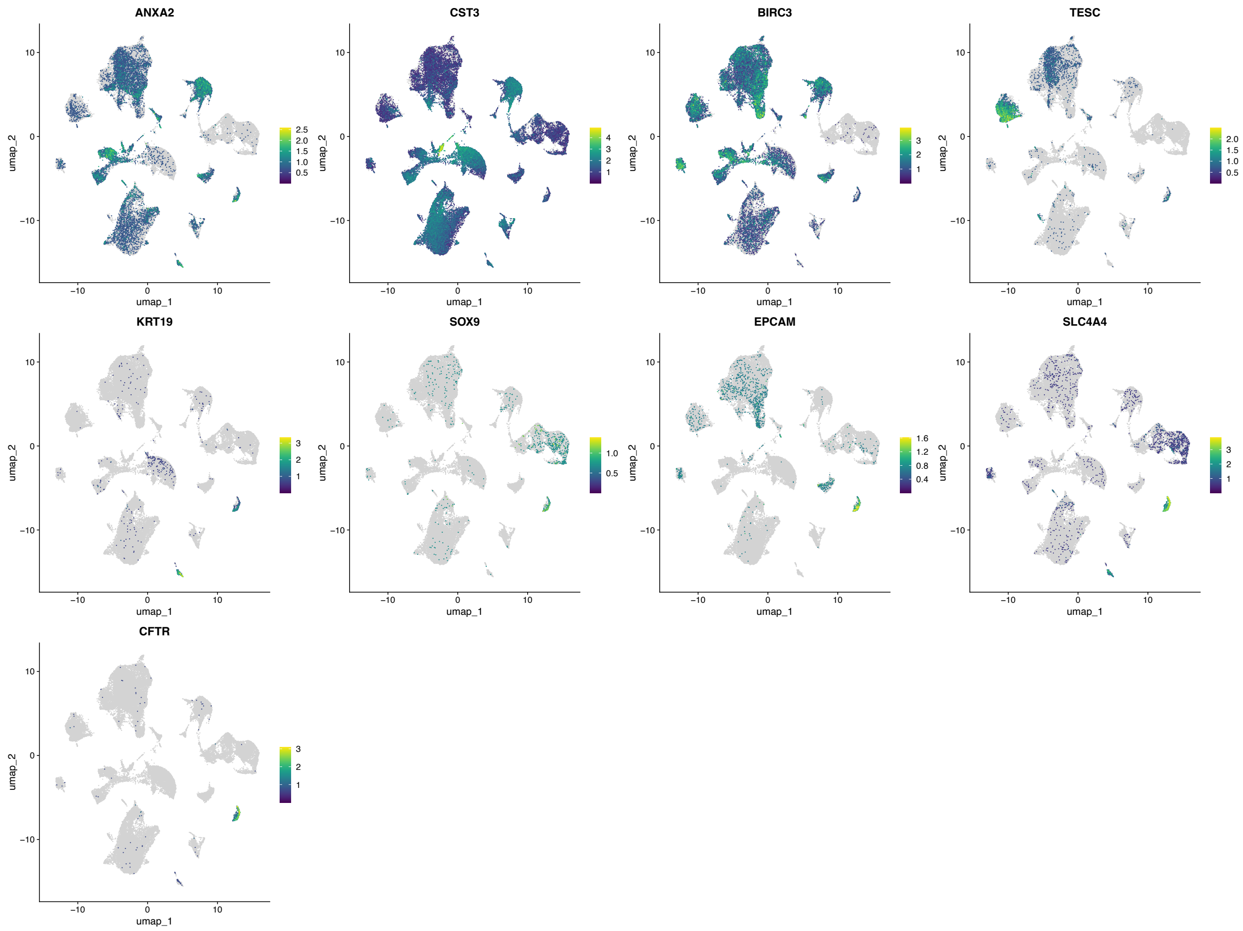

**Supplementary figure 7. Cholangiocyte markers in human distinguish small populations in the woodchuck liver.** Marker genes found in human cholangiocytes expressed in woodchucks: *ANXA2, CST3, BIRC3, TESC, KRT19, SOX9* and *EPCAM.*

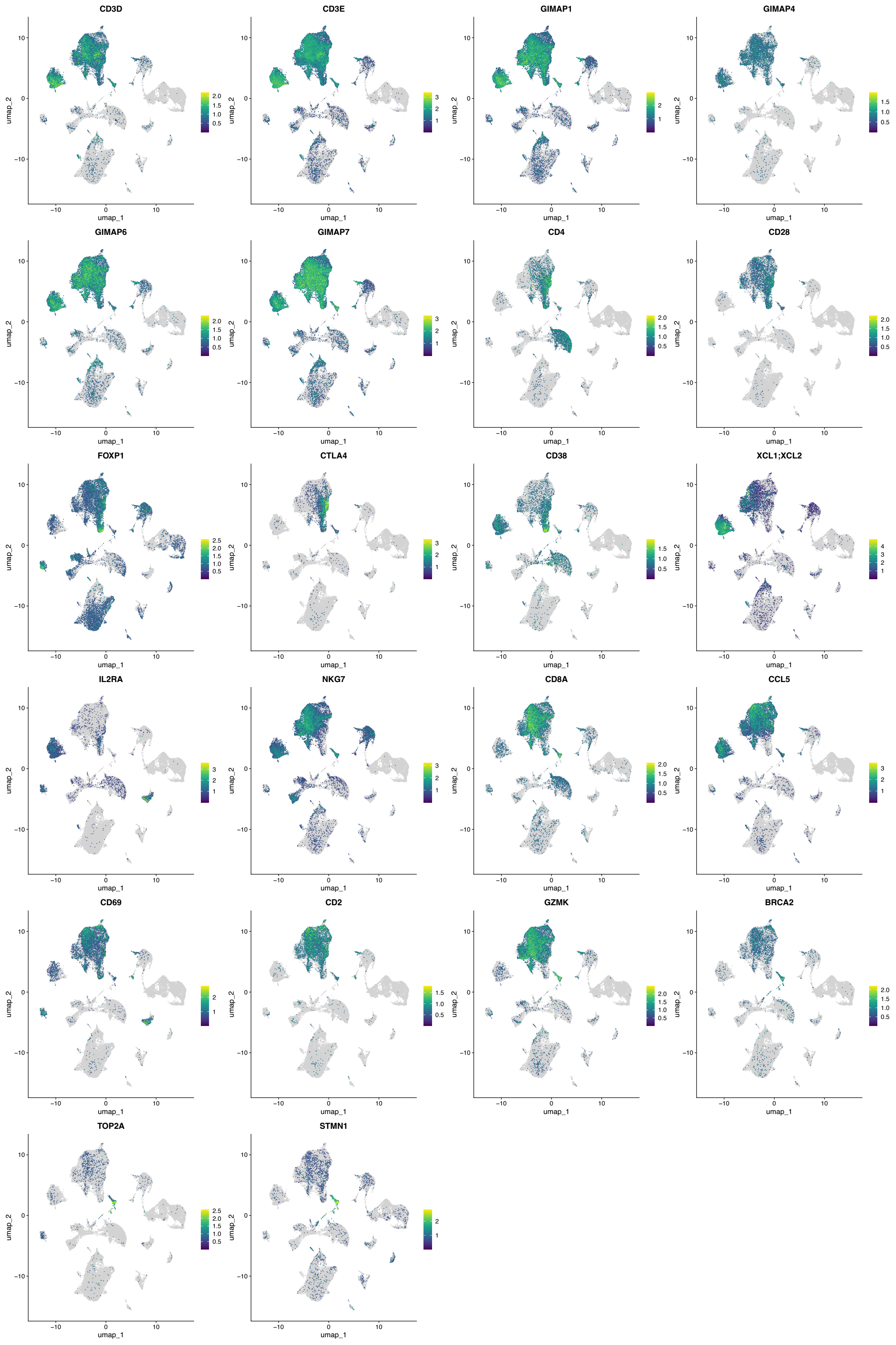
CD3D
CD3E
GIMAP1
GIMAP4
GIMAP6
GIMAP7
CD4
CD28
FOXP1
CTLA4
CD38
XCL1;XCL2
IL2RA
NKG7
CD8A
CCL5
CD69
CD2
GZMK
BRCA2
TOP2A
STMN1
umap_1
umap_2

**Supplementary figure 8. Woodchuck T cell subsets in the liver can be distinguished by human markers.** Significant genes distinguishing T lymphocytes and lymphocyte subtypes in the liver. Gene expression values were projected as heatmaps onto a UMAP of woodchuck liver cells. Markers distinguishing T lymphocytes include *CD3D, CD3E, GIMAP1, GIMAP4, GIMAP6*, and *GIMAP7*. Naïve regulatory T cells express: *CD4, CD28, FOXP1, CTLA4*, and *CD38*. NK/T cells express: *XCL1; XCL2, IL2RA (CD25),* and *NKG7*. Tissue-resident memory T cells express: *CD8A, CCL5, CD69, CD2, GZMK* but not *CD25* or *CD38*. Proliferating T cells express *BRCA2, TOP2A*, and *STMN1.*

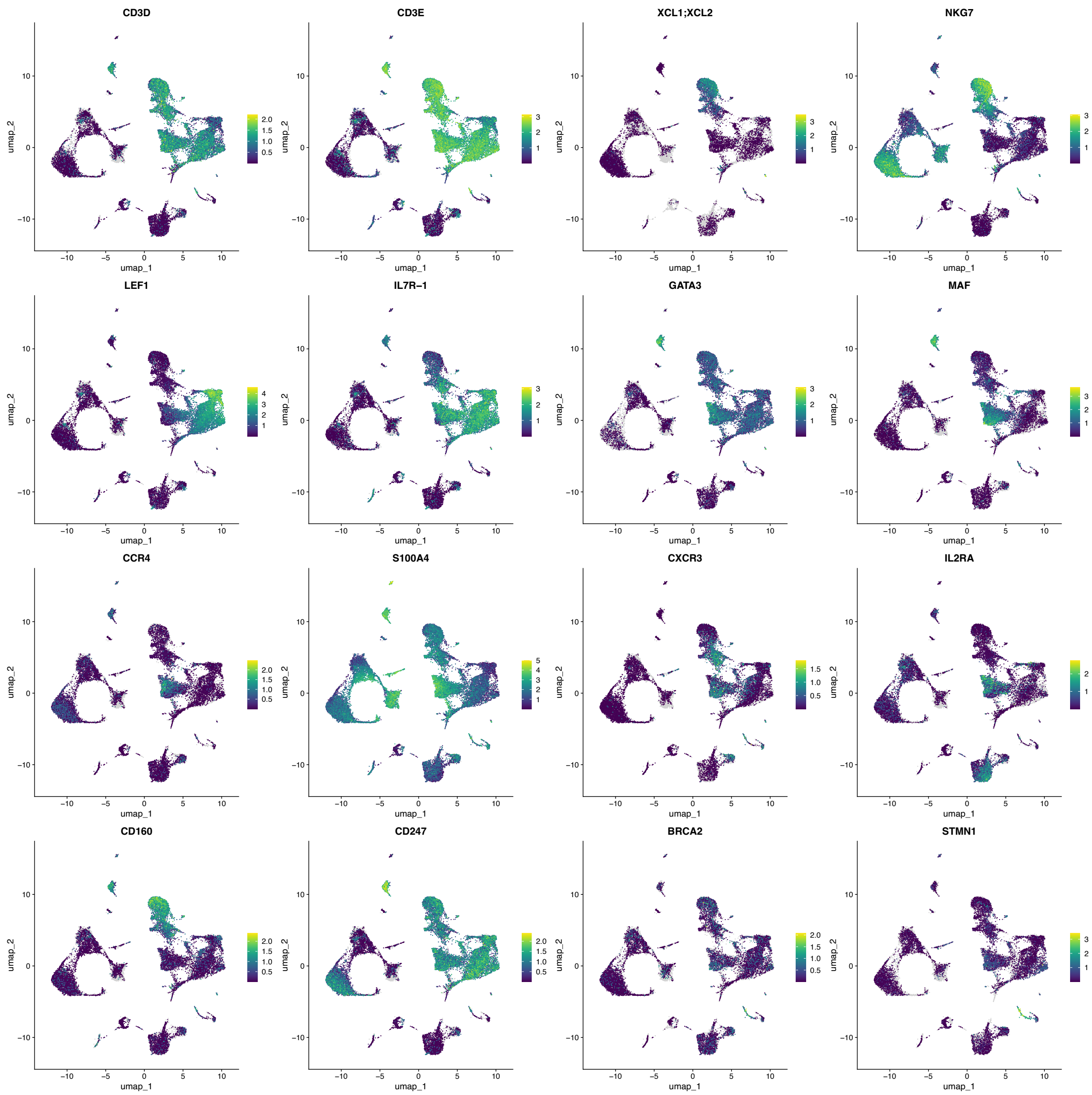

**Supplementary figure 9: T-cell subsets in woodchuck PBMCs can be distinguished by human markers.** Important genes distinguishing T lymphocytes and lymphocyte subtypes in PBMCs. Gene expression values are projected as heatmaps onto a UMAP of woodchuck PBMCs. Markers distinguishing all T lymphocytes include *CD3D* and *CD3E*. NK/T cells express: *XCL1; XCL2,* and *NKG7*. Memory T cells express: *LEF1* and *IL7R*. TH2-like T cells 1 and 2 express: *GATA3, MAF, CCR4*, and *S100A4*, whereas population 1 more strongly expresses *CXCR3* and *IL2RA (CD25)*, and population 2 expresses *CD160*. Proliferating T cells express *BRCA2* and *STMN1*.

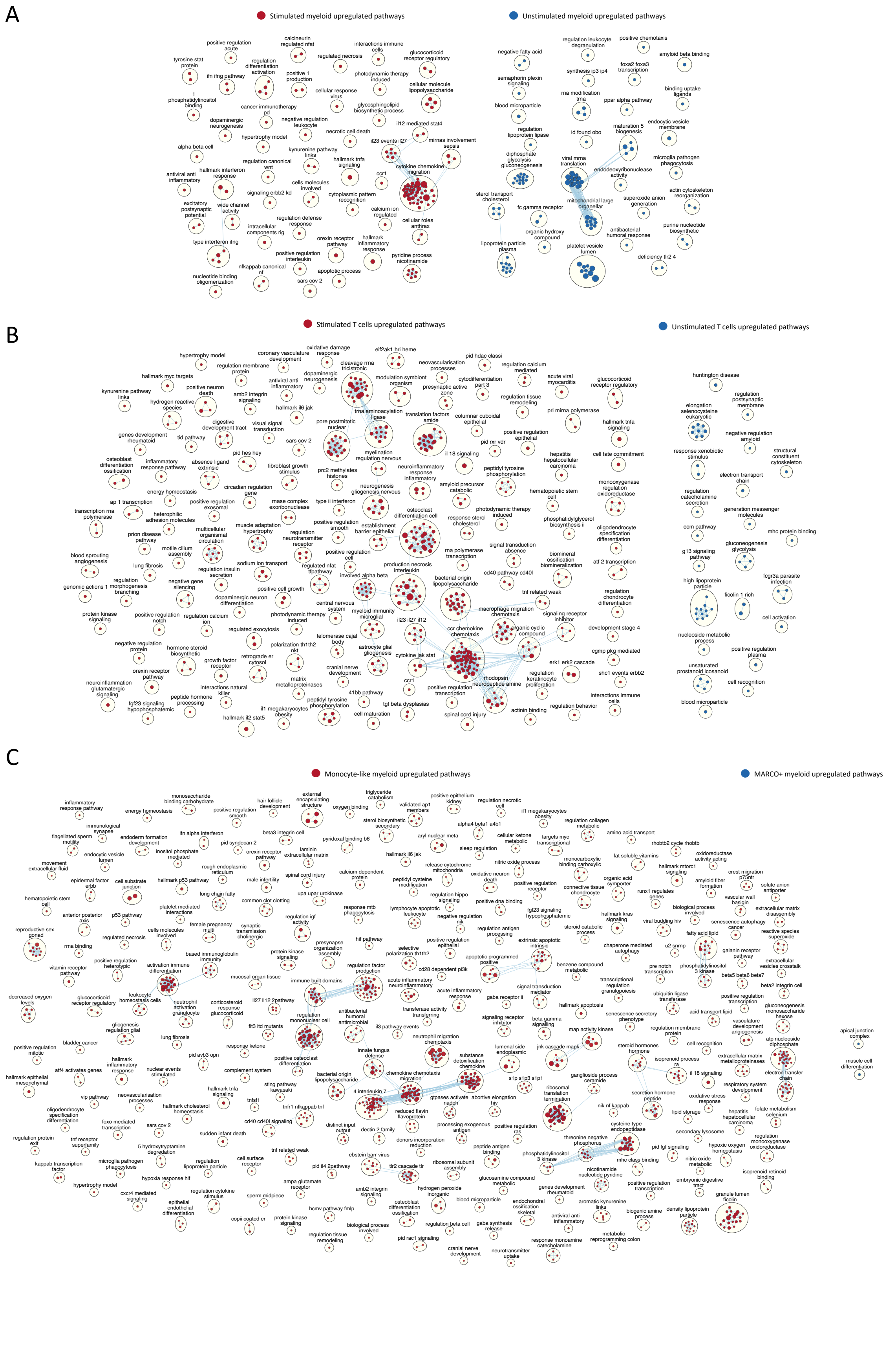
A
Stimulated myeloid upregulated pathways
Unstimulated myeloid upregulated pathways
B
Stimulated T cells upregulated pathways
Unstimulated T cells upregulated pathways
C
Monocyte-like myeloid upregulated pathways
MARCO+ myeloid upregulated pathways

**Supplementary figure 10: Pathway analysis of myeloid and T cells in the stimulated data.** Pathway enrichment analysis was performed on DE genes found in the stimulated dataset when directly comparing two populations to each other. Pathways were identified with GSEA and visualized with Cytoscape. Each dot (node) represents a pathway, with the size of the dot indicating the number of genes in the pathway. Related pathways are connected by light blue lines, grouped together in black circles, and labelled. (A) Pathways associated with stimulated myeloid cells (red) and unstimulated myeloid cells (blue) from a direct population comparison. (B) Pathways associated with stimulated T cells (red) and unstimulated T cells (blue). (C) Pathways associated with monocyte-like myeloid cells (red) and MARCO+ myeloid cells (blue) from both the stimulated and unstimulated datasets.

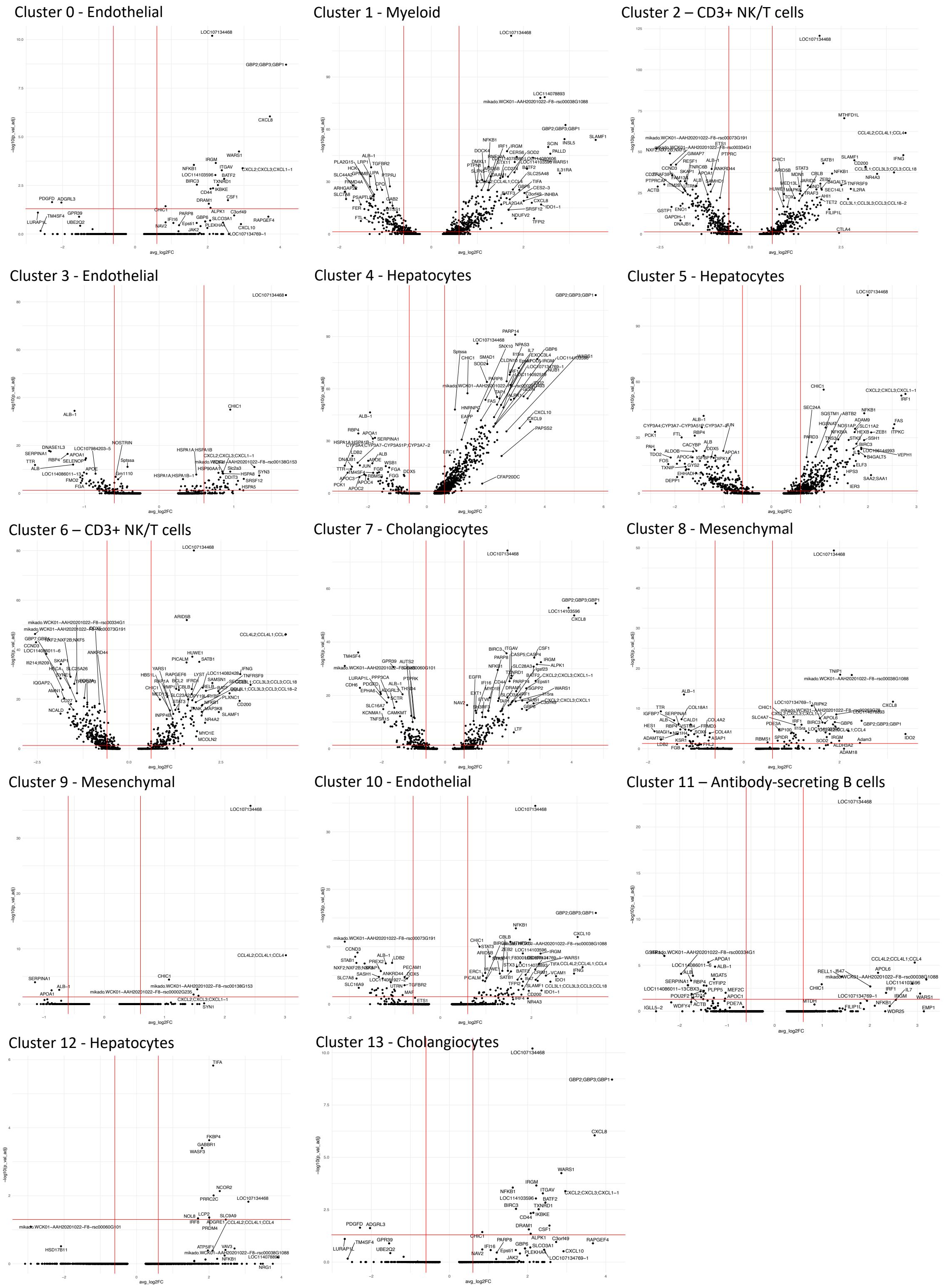

**Supplementary figure 11: Volcano plots of stimulated versus unstimulated data.** Volcano plots indicating key differentially expressed genes in cell clusters (as seen in

Fig. 8) present in control and PMA/ionomycin stimulated precision cut liver slice woodchuck tissue.

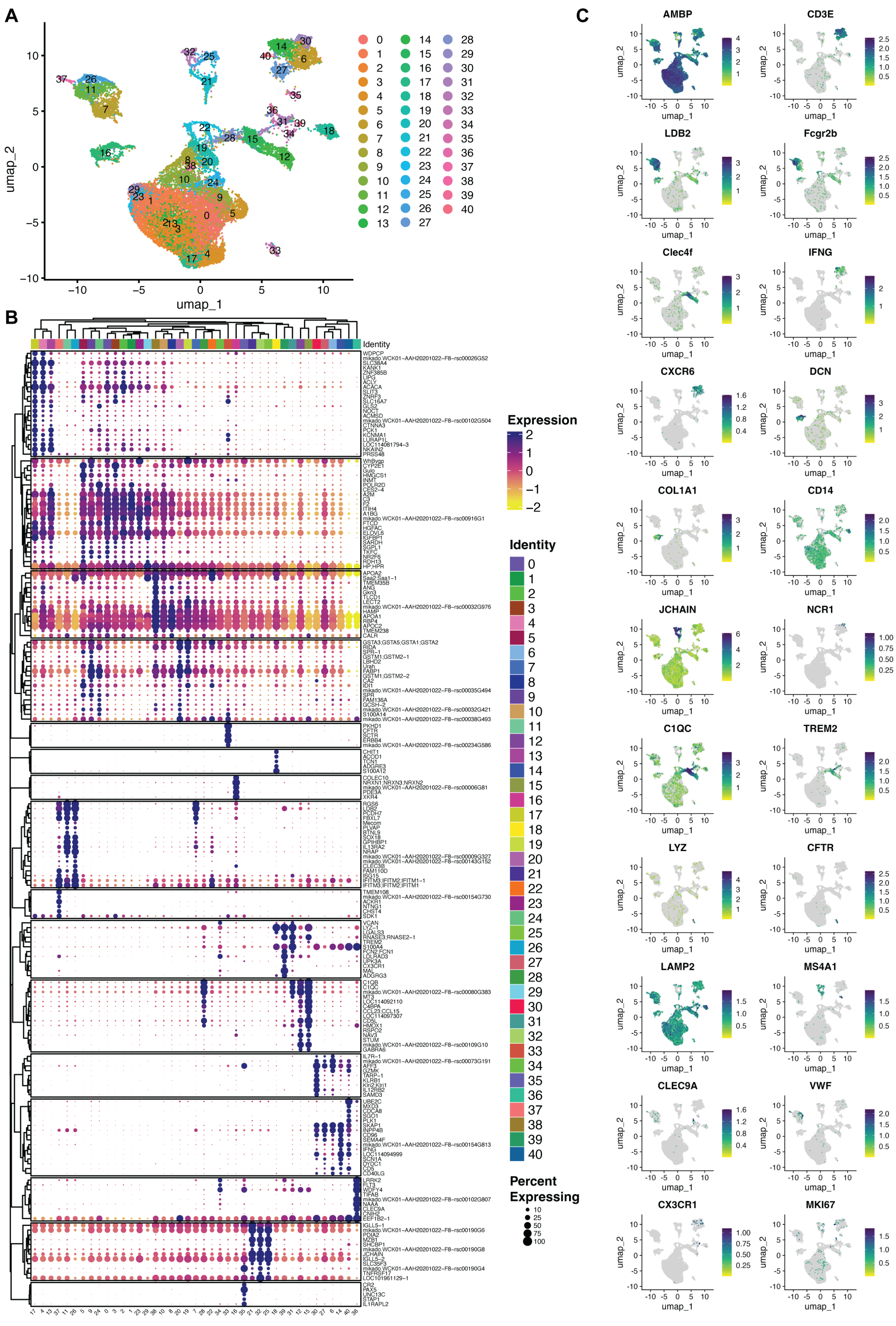
A
B
C
umap_1
umap_2
Identity
Expression
Percent Expressing
AMBP
CD3E
LDB2
Fcgr2b
Clec4f
IFNG
CXCR6
DCN
COL1A1
CD14
JCHAIN
NCR1
C1QC
TREM2
LYZ
CFTR
LAMP2
MS4A1
CLEC9A
VWF
CX3CR1
MKI67

**Supplementary figure 12: WHV dataset analysis workflow.**
(A) UMAP indicating healthy and chronically infected WHV carrier data with 41 clusters.
(B) Clustered heatmap indicating the top differentially expressed gene per cluster in (A).
(C) Feature UMAPs displaying the expression of key cell-type specific marker genes.

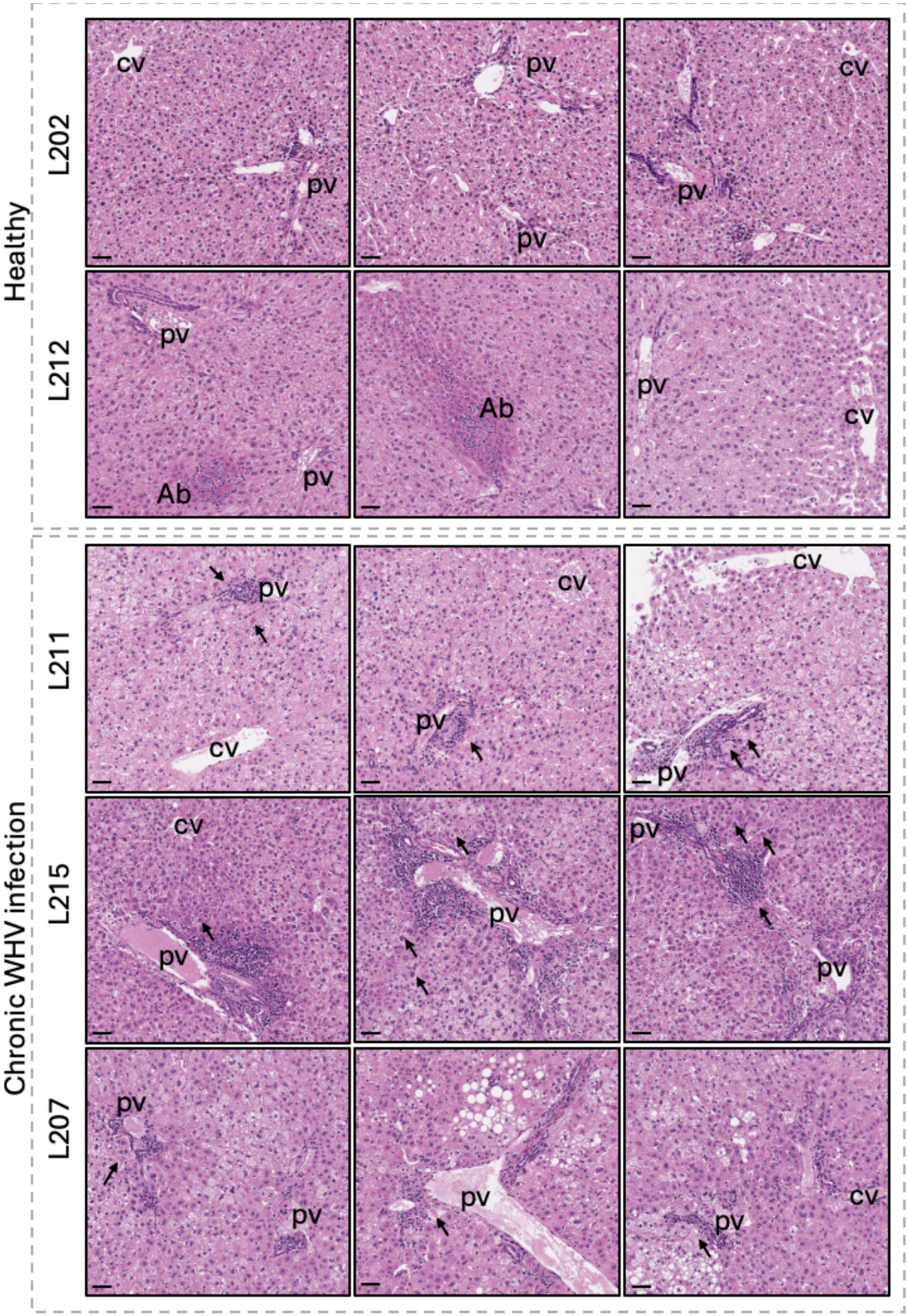
Healthy
L202
cv
pv
pv
pv
cv
pv
L212
pv
Ab
pv
Ab
pv
cv
Chronic WHV infection
L211
pv
cv
cv
pv
cv
pv
L215
cv
pv
pv
pv
pv
L207
pv
pv
pv
pv
cv

**Supplementary figure 13: Histological evaluation of healthy and cWHV woodchuck tissue using H&E.**

Three representative images were selected for each animal. Central veins (cv) and portal veins (pv) are labelled with text while areas of hepatocyte necrosis are indicated by arrows. Chronically infected woodchuck livers display a moderate hepatitis: grade 2-3 /4 of activity scoring. Infiltration of cells and hepatocyte necrosis is localised to periportal regions while central veins are spared. There are no indications of bridging necrosis or multi/pan-acinar necrosis within cWHV tissue. Within healthy sample L212, interzonal hepatocytic micro-abscesses infiltrated with neutrophils are present as a result of acute reperfusion ischemic injury. Scale bars represent 50μm.

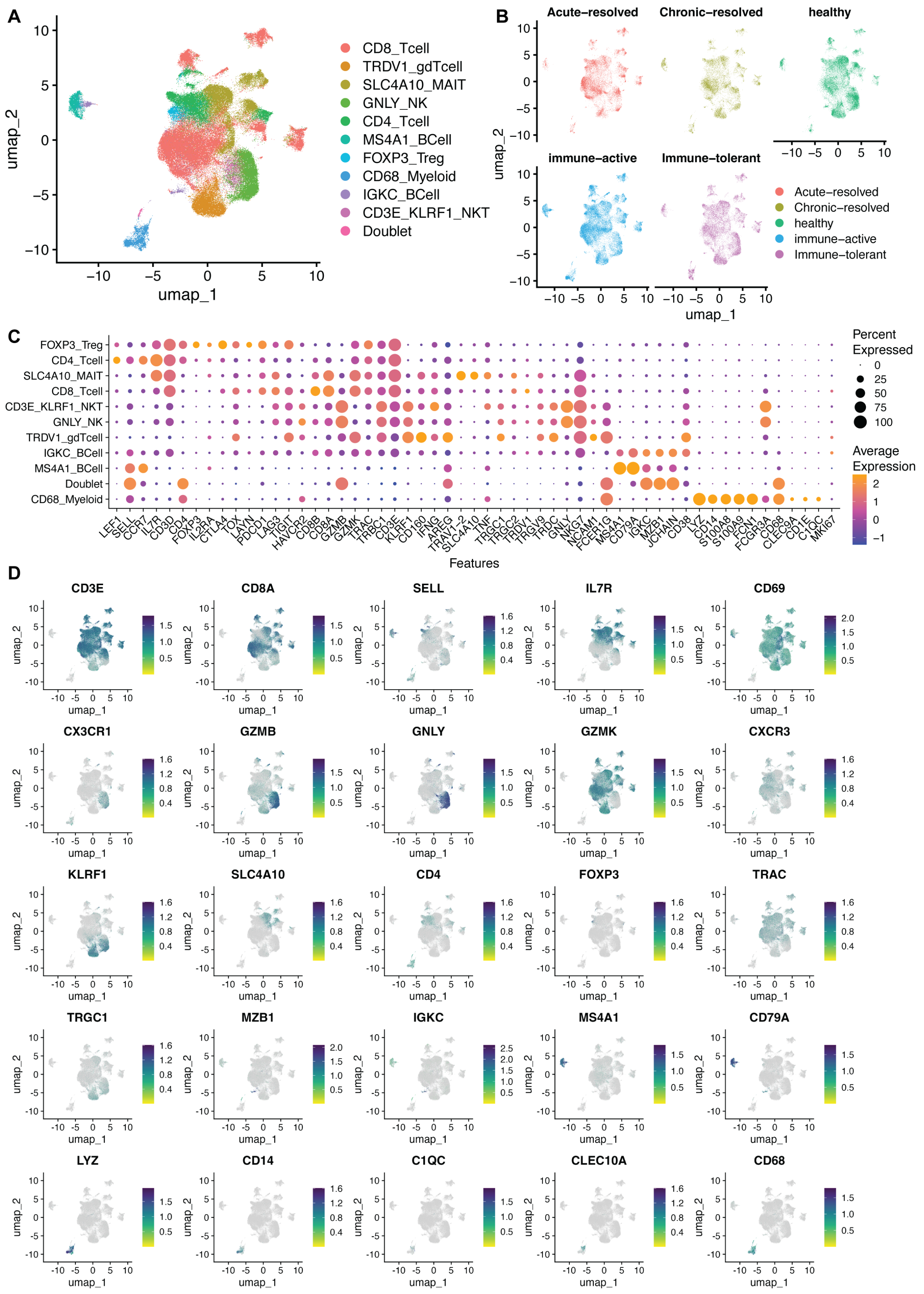
A
B
C
D
CD8_Tcell
TRDV1_gdTcell
SLC4A10_MAIT
GNLY_NK
CD4_Tcell
MS4A1_BCell
FOXP3_Treg
CD68_Myeloid
IGKC_BCell
CD3E_KLRF1_NKT
Doublet
umap_1
umap_2
Acute−resolved
Chronic−resolved
healthy
immune−active
Immune−tolerant
Percent Expressed
Average Expression
Features
CD3E
CD8A
SELL
IL7R
CD69
CX3CR1
GZMB
GNLY
GZMK
CXCR3
KLRF1
SLC4A10
CD4
FOXP3
TRAC
TRGC1
MZB1
IGKC
MS4A1
CD79A
LYZ
CD14
C1QC
CLEC10A
CD68

**Supplementary figure 14: HBV dataset annotation and preparation for comparison to cWHV samples.**
A. Cell-type annotated UMAP of core-needle biopsy samples from B. acute-resolved, chronic-resolved, healthy, immune-active and immune-tolerant human samples (GSE182159). C. Expression of lineage-associated marker genes per cluster to support cluster annotations. D. Feature UMAPs indicating the expression of key lineage-associated marker genes.

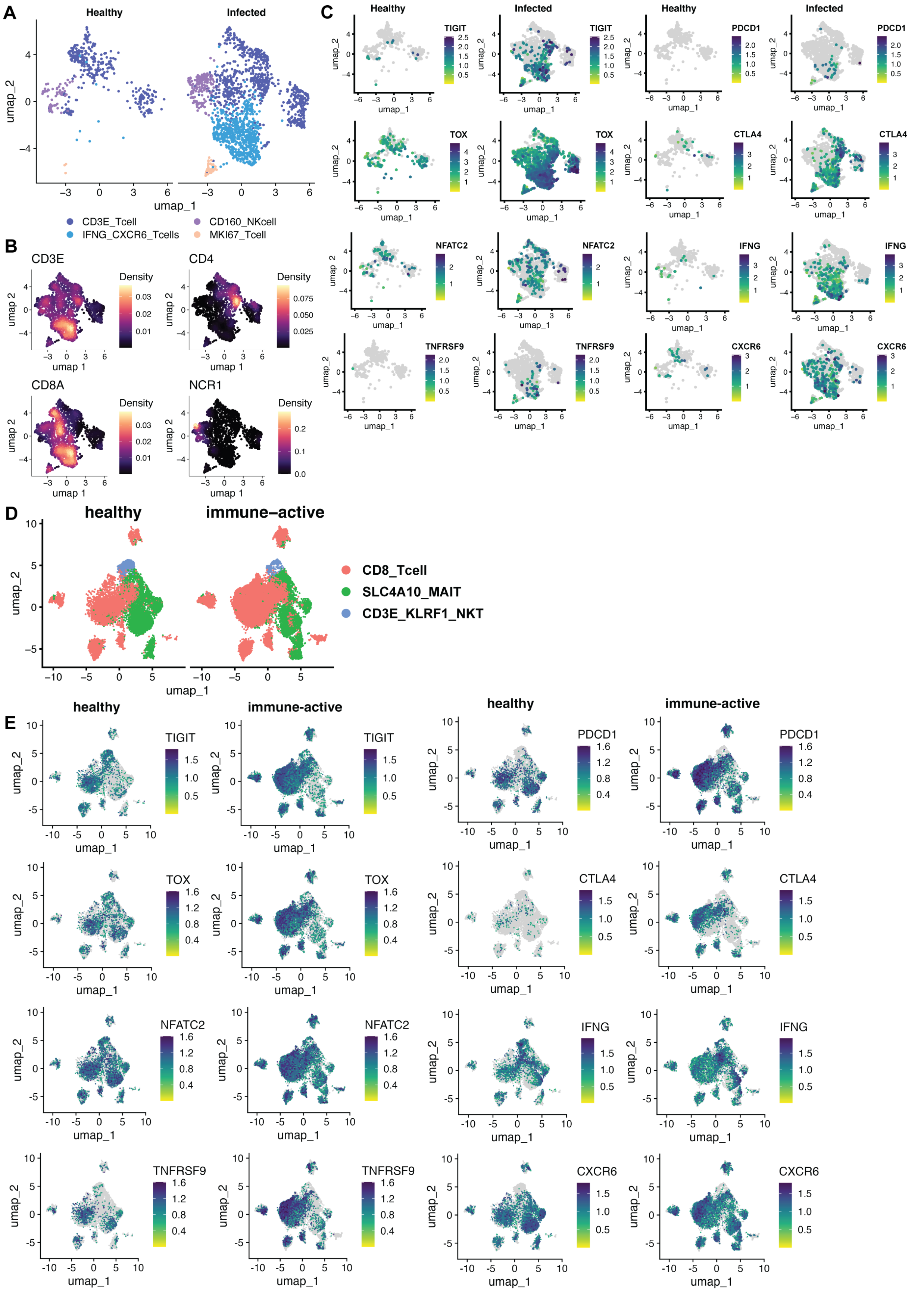
A
Healthy
Infected
umap_2
umap_1
CD3E_Tcell
IFNG_CXCR6_Tcells
CD160_NKcell
MKI67_Tcell
B
CD3E
CD4
CD8A
NCR1
Density
umap 1
umap 2
C
Healthy
Infected
TIGIT
PDCD1
TOX
CTLA4
NFATC2
IFNG
TNFRSF9
CXCR6
D
healthy
immune–active
CD8_Tcell
SLC4A10_MAIT
CD3E_KLRF1_NKT
E
healthy
immune-active
TIGIT
PDCD1
TOX
CTLA4
NFATC2
IFNG
TNFRSF9
CXCR6

**Supplementary figure 15: Expression of select genes in T cells shared between WHV and HBV chronic infections.**

A. Sub-clustering of lymphocytes from healthy and cWHV samples. B. Expression of T cell *(CD3E, CD4, CD8A)* and NK-cell *(NCR1)* specific genes in the sub-clustered map. C. Expression of T cell activation and exhaustion associated genes in the cWHV T cell sub-clustered map. D. Sub-clustered map of *CD8A* expressing conventional and innate T cell populations from healthy and immune-active chronically infected HBV core-needle biopsy samples. E. Expression of T cell activation and exhaustion associated genes in the HBV infected T cell sub-clustered map.

**Supplementary table legends**

**Supplementary table 1: CellRanger and cell type statistics per woodchuck sample**

**Supplementary table 2: DE genes for cell types in the woodchuck liver.** DE genes were calculated using an implementation of Presto run within Seurat (`RunPrestoAll`).

**Supplementary table 3: DE genes for clusters in the woodchuck liver.** DE genes calculated on a high-resolution clustering rather than based on their general cell type label using Presto.

**Supplementary table 4: DE genes for cell types in woodchuck PBMCs.** DE genes calculated with Presto based on cell type.

**Supplementary table 5: DE genes for clusters in woodchuck PBMCs.** DE genes calculated with Presto based on high resolution clustering.

**Supplementary table 6: DE genes for spatial transcriptomics clusters.** DE genes calculated using the Seurat function `FindAllMarkers`.

**Supplementary table 7: List of DE genes in a periportal vs pericentral comparison from spatial transcriptomics data ranked by Wstat.**

**Supplementary table 8: List of genes correlated to periportal and pericentral gene signatures.** Gene counts from SCT slot in Seurat object correlated to gene signatures with Spearman correlation.

**Supplementary table 9: DE genes for clusters in stimulated dataset.** DE genes calculated using the Seurat function `FindAllMarkers` at a cluster-level resolution.

**Supplementary table 10: DE genes for a comparison of stimulated vs unstimulated cells for each cluster in the stimulated datasets.** DE genes calculated with Presto.

**Supplementary table 11: DE genes for clusters in Healthy+WHV-infected Woodchuck map.** Calculated using Presto.

**Supplementary table 12: Pairwise disease vs healthy DE genes per cell type present in the Healthy+WHV-infected woodchuck map.** Calculated using Seurat's FindMarkers (log.FC=0.1, min.pct=0.15, only.pos=TRUE, test.use="wilcox")

**Supplementary table 13: Cytotoxic and exhaustion associated gene signatures used to generate Figure 5G.**

**Supplementary table 14: Paired orthologs for woodchuck genes.** Genes found using pairwise ortholog results between woodchuck and all other species. One-to-many orthologs are connected with semi-colons. NAs represent no ortholog found in that species. “ybmar”, “almar”, and “squir” represent the yellow-bellied marmot, Alpine marmot, and 13-lined ground squirrel respectively. “hierGene” lists the ortholog used to label the woodchuck gene, and “uniqueHier” is a unique version of that gene (i.e. with duplicates labeled with a -1, -2, etc. added at the end of the gene symbol). “humanOneToOne” and “mouseOneToOne” are one-to-one orthologs between woodchuck and human and mouse respectively.

**Supplementary table 15: Human and mouse genes not found in the woodchuck genome**
